# Effect of cell membrane tension on the lifetime and size of mature clathrin-coated pits and their spatial distribution

**DOI:** 10.1101/2023.03.16.532501

**Authors:** Xinyue Liu, Wang Xi, Xiaobo Gong

**Affiliations:** Shanghai Institute of Applied Mathematics and Mechanics, School of Mechanics and Engineering Science, Shanghai Frontier Science Center of Mechanoinformatics, Shanghai Key Laboratory of Mechanics in Energy Engineering, Shanghai University, Shanghai 200072, China; Mechanobiology Institute, National University of Singapore, 5A Engineering Drive 1, Singapore, 117411, Singapore; Institut Jacques Monod, Université Paris Cité, Centre National de la Recherche Scientifique, Paris F-75013, France; Key Laboratory of Hydrodynamics (Ministry of Education), Department of Engineering Mechanics, School of Naval Architecture, Ocean and Civil Engineering, Shanghai Jiao Tong University, Shanghai 200240, China; Institute of Mechanobiology and Biomedical Engineering, School of Life Science and Technology, Shanghai Jiao Tong University, Shanghai 200240, China

**Keywords:** Clathrin-coated pit, CCP lifetime, CCP size distribution, mature CCP size, membrane tension

## Abstract

Clathrin-mediated endocytosis is the most characterized pathway for cells to internalize diverse receptor-bound cargo, such as proteins, nanoparticles, and viruses. However, the effect of membrane tension on clathrin-coated pit (CCP) maturation remains inadequately characterized. This study aimed to determine the effect of membrane tension on CCP maturation both spatially and temporarily, which remains a controversial and elusive issue. We obtained the sizes and spatial distributions of CCPs by the structured illumination microscopy of fixed cells and observed CCP lifetimes in live cells by total internal reflection fluorescence microscopy. The processes of CCP maturation and abortion were reproduced numerically through Monte Carlo simulation. The results showed that the growth time of CCP was more reasonably proportional to its volume rather than its surface area. We further investigated the spatial distribution of the membrane tension and size of CCPs, finding a significant positive correlation between the membrane tension and the size of mature CCPs spatially. This indicates that the CCPs tend to enrich in the highest-tension region, especially the mature ones. These results agreed with our numerical prediction that the CCP structure grew larger to overcome a higher energy barrier caused by higher background cell membrane tension. Our findings enhance the understanding of CCP maturation dynamics and underscore the importance of membrane tension in regulating CCP development.

## INTRODUCTION

Clathrin-mediated endocytosis is a widely studied process in vesicular trafficking that transports cargo molecules from the cell surface to the interior [1]. Increased membrane tension inhibits endocytosis [2] on the subcellular scale by counteracting clathrin polymerization [3] and membrane fission [4]. However, understanding the lifetime and size of clathrin-coated pits (CCPs) affected by the membrane tension remains elusive.

The size and lifetime of CCP are two factors highly correlated with their productivity. Two typical CCP catalogs were characterized through a global analysis of total internal reflection fluorescence (TIRF) microscopy time series [5]: one was classified as productive that takes 30–120 s to mature, and the other as abortive, with typical lifetimes (time from formation to extinction) of less than 20 s. It is believed that endocytic cargo plays a crucial role in deciding the fate of a CCP [6], and cargo binding stabilizes the CCPs and facilitates their growth toward vesicle formation. Banerjee et al. [7] developed a coarse-grained model of CCP by mapping the CCP assembly dynamics onto a one-dimensional random walk to explain the relationship between the CCP size and lifetime. Using Monte Carlo simulations, they analytically examined the statistical properties of the lifetimes and predicted the maximum size of abortive CCPs to be around 90 nm [8]. CCP structures are of various curvatures and sizes, ranging from 60–120 nm [9, 10], which is below the 200-nm diffraction limit of fluorescent microscopes such as TIRF or confocal microscopes. In practical terms, only cellular structures and objects at least 200–350 nm apart can be resolved [11]. Li et al. [12] extended the resolution of live-cell structured illumination microscopy (SIM) to 45–62 nm and revealed a positive correlation of 0.20 nm/s between CCP diameter and growth time by linear regression. Willy et al. [13] observed de novo CCPs using SIM and found that they developed a curvature in the early stages of their formation. However, to date, the effect of cell membrane tension on CCP size and lifetime still cannot be investigated directly.

More importantly, seminal studies conducted at the beginning of the last decade have demonstrated that cell mechanics, in particular the changes in plasma membrane tension, give rise to dynamically distinct populations of clathrin-coated structures within cells [14]. It has been suggested that tension-dependent endocytosis and exocytosis are involved in surface area regulation and buffering of membrane tension. High membrane tension leads to excess exocytosis, an increase in cell surface area, and a decrease in tension, and vice versa [15]. Overall, the endocytic vesicle formation is slowed down by increased membrane tension, as high tension resists the generation of curved clathrin coats [3, 16, 17]. In a majority of these studies, the dynamics of CCSs were monitored at the plasma membrane–coverglass interface; it was found that the cell spreading area could regulate clathrin dynamics by qualitatively controlling the tension of the membrane in vitro [18, 19]. In practice, the local cell membrane tension on the adherent surface is difficult to be measured directly or predicted accurately with membrane–substrate interactions, confined by focal adhesions. Therefore, the observation of the bottom surface of the cell alone does not give a global picture of this problem.

Crucially, several studies have elucidated the role of membrane tension in CCP dynamics. The role of specific proteins in mediating the effects of membrane tension on CCP formation has been highlighted. Willy et al. [20] emphasized the importance of clathrin assembly lymphoid myeloid leukemia protein (CALM) in supporting CCP completion under increased membrane tension. Meanwhile, Willy et al. [21] explored the spatiotemporal heterogeneity of CCP dynamics influenced by membrane mechanics, shedding light on the variability of CCP behavior across different cellular regions. Batchelder and Yarar [22] found that cell-substrate adhesion reduced the rate of CCPs formation and with a stronger requirement for actin polymerization in areas of adhesion. Research by Akatay et al. [23] provided evidence for the formation of giant CCPs under elevated membrane tension, with actin dynamics playing a critical role in this process. Gauthier et al. [24] demonstrated that membrane tension decreases during cell spreading, facilitating the incorporation of new membrane areas through exocytosis. Despite these advancements, significant controversies remain. The influence of substrate stiffness on membrane tension, as debated by Kreysing et al. [25], suggests that effective cell membrane tension remains largely unaffected by changes in the stiffness of the polyacrylamide substrate, thereby not affecting endocytosis in the manner previously thought. Furthermore, the exact molecular mechanisms by which proteins like CALM [26, 27], adaptor protein (AP2) [28, 29] and actin [30] dynamics facilitate CCP formation under varying membrane tensions are still not fully understood.

The formation of CCPs is highly energetically demanding, particularly under conditions of elevated membrane tension, which necessitates additional mechanisms to facilitate the necessary membrane curvature. Wu and Wu [31] discussed the coupling of clathrin turnover with cargo sorting, indicating that ATP hydrolysis by Hsc70 and auxilin is necessary for efficient clathrin uncoating and cargo selection. They mentioned that clathrin assembly can be reconstituted in vitro without external energy, but the coupling of cargo enrichment and clathrin turnover requires energy support. Sochacki et al. [32] proposed a model where CCP curvature is activated by the release of a flattening force, enabling the triskelia network to spontaneously transition into a more energetically favorable state. Saleem et al. [3] demonstrated that elevated membrane tension increases the energy cost of curvature generation, highlighting the need for additional force or energy input to facilitate this process. Zeno et al. [33] revealed that clathrin does not merely passively assemble at sites of high curvature but actively senses and prefers these regions. Tagiltsev et al. [34] proposed a new energy landscape model for CCPs, suggesting that clathrin lattices with curvatures less than the intrinsic curvature of clathrin triskelia are elastically loaded, storing elastic energy and drive the lattice to bend into highly curved CCPs. The research highlights the energy contributions to CCP formation, including the energy required to bend the cell membrane, lateral tension, the bending energy of the clathrin coat, and the energy of clathrin polymerization. Aguet et al. [28] suggested that curvature generation within nascent CCPs is a critical factor for their maturation and that CCPs that fail to gain curvature are aborted. However, the energy insights into the CCP maturation process without requiring detailed assumptions about the temporal evolution of CCPs, which are challenging to observe experimentally.

In this study, we focus on two scientific questions: 1) the relationship between CCP lifetime and maturation size, specifically whether we can obtain a curve depicting the changes in CCP size over time by separately observing the spatial distribution of CCP sizes and their lifetimes; and 2) how background membrane tension affects the maturation size and lifetime of CCPs before their pinch-off.

To address the first question, we propose a method that combines TIRF and SIM observations to obtain the curve of CCP size and lifetime. We measured the sizes and locations of CCPs using 3D scans from SIM and acquired the lifetimes of CCPs using TIRF. TIRF is widely used in many studies of CCPs [5,35,36] due to its high temporal resolution and less laser exposure, which minimizes fluorescence quenching and allows prolonged observation of cells. However, the disadvantage of TIRF is that it can only observe the cell bottom, based on the total reflection principle of light [37]. The spatial resolution of TIRF is also insufficient to observe the size changes of CCPs during their formation. Although SIM offers significantly improved spatial resolution compared to TIRF, it is successful for individual CCP measurements [13] but challenging for high-throughput measurements of CCPs across an entire cell. SIM takes approximately 30 minutes to scan a whole cell, during which it cannot capture clear images due to the migration of living cells. Consequently, cells must be fixed during SIM scanning, preventing direct observation of CCPs lifetimes. Despite these challenges, we aimed to complement SIM and TIRF observations. First, we reproduced the growth process of CCPs observed using TIRF through Monte Carlo simulations and examined two possible relationships between CCP size and lifetime by comparing the CCP size distribution from the simulations with results observed using SIM. Then, we determined the size ranges of de novo, developing, and mature CCPs based on the relationship between CCP size and lifetime.

To address the second question, we extracted the cell shape from the spatial distribution of CCPs observed by SIM. We simulated the apparent membrane tension distribution of a free-spread cell in arbitrary shape using an algorithm proposed by our previous study [38]. The apparent membrane tension was then considered as the background membrane tension of CPPs locations. Further investigation of the spatial distribution of membrane tension and CCP size revealed a significant positive correlation between the membrane tension and the CCP size spatially, indicating that the CCPs tended to enrich in the highest-tension region, especially the mature ones. Additionally, the CCP size distribution showed heavy-tailed compared to the standard normal distribution, leading us to explore whether it was caused by membrane tension distribution. It has been reported that endocytosis is randomly initiated and then stabilized by CCPs [36], and our previous study [39] supported the idea that the CCPs act as a mechanical reinforcement in the wrapping region, helping to fend off external forces during endocytosis. We found that larger CCP structures are needed to overcome a higher background membrane tension to assist the cargo uptake. In the temporal aspect, the energy barrier caused by the increase in tension leads to a decrease in the number of de novo CCPs [19] and an increase in the lifetime of larger-sized CCPs [12]. Interestingly, in the spatial aspect, the large-sized and long-lived CCPs were revealed to concentrate in the high-tension region in this study.

## RESULTS

### CCP size and distribution on the cell membrane

To obtain the statistical properties of the CCPs spatial distribution, a total of 12 spread RBL-2H3 cells were scanned using SIM. Figure 1 A shows the reconstructed fluorescence image of Cell ID. 09. The structured light reconstruction images were obtained using Nikon imaging software with a resolution of about 50 nm [40, 41]. The method of Gaussian kernel fitting to identify particles [42] is used here to compensate for the spatial resolution. The sizes and positions of CCPs were identified using Imaris software, and the CCPs on the cell surface were extracted using custom MATLAB code. The 3D numerical reconstruction of the cell structure was performed using the built-in library function of MATLAB for convex or concave hull boundary searching based on the positions of CCPs. The geometric parameters, including cell volume, cell area, and spread area, are listed in Table S1 (Supporting Information). Heuser et al. [9] observed using the cryo-electron microscope that the size range of CCPs was 60–120 nm. It is important to note that the CCP size observed using the cryo-electron microscope is close to its actual size, whereas the size observed using fluorescence microscope appears much larger. This discrepancy occurs because the fluorescence microscopy is limited by the diffraction of light, causing the observed size of small particles to appear larger due to the point spread function (PSF) of the microscope. For example, in the SIM observation of Li et al. [12], the CCPs were large and stable enough to be resolved as a ring, which finally grew to a median maximum diameter of 152 nm. Therefore, the CCPs sizes mentioned in this study are regarded as SIM observation size. In addition, the resolutions of the Z direction and the XY plane differ for a Gaussian laser beamm, so the obtained CCP diameters were considered equivalent diameters.

**Figure 1:**
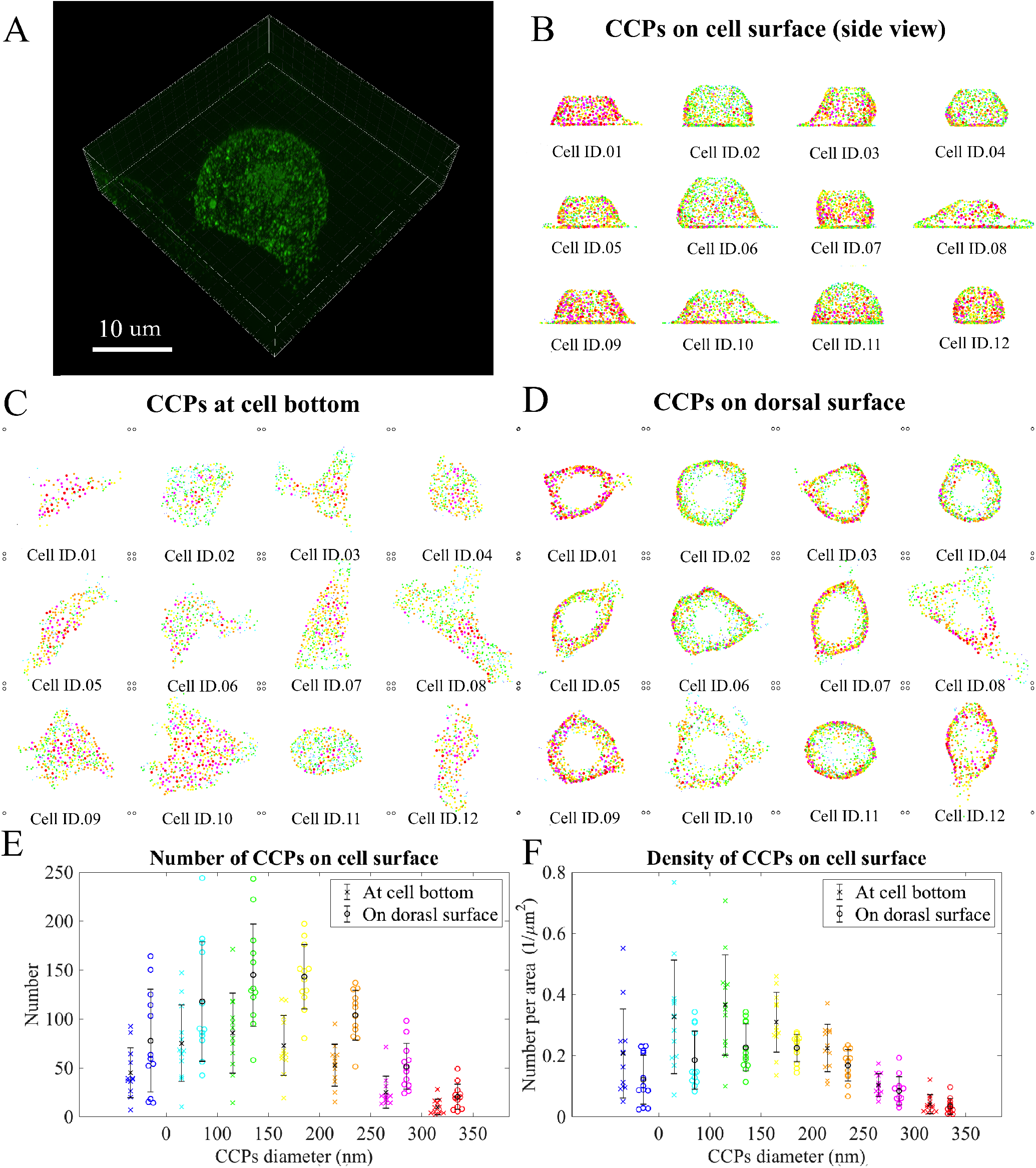
CCP spacial distribution is obtained by SIM. (A) A fluorescent image of CCPs on Cell ID.09 was chosen as a demonstration. The CCP locations were identified (B) on the total surface (side view), (C) on the bottom surface (bottom view), and (D) on the dorsal surface (top view) in the boxes of 32 *×* 32 *×* 15 *µ*m^3^. The distributions of CCP number (E) and density (F) were sorted by size on a total of 12 scanned cells.

Figure 1 B shows the side view of the positions and sizes of CCPs. The CCP sizes were divided into seven ranges, represented by seven different colors. The CCP sizes represented by each color in all subsequent figures are consistent with those in this figure. The CCP positions and sizes on the bottom surface appear randomly distributed to the naked eye (Figure 1 C); interestingly, the CCPs are rarely observed at the center of the cell top (Figure 1 BD). Consistently, Grossier et al. [43] found that the endocytosis of Tf occurred mostly near the bottom of the cell, with almost no endocytosis on the top surface of the cell. Similarly, some epidermal growth factor endocytosis occurred on the bottom surface of the cell, was evenly distributed on the surface of the middle part of the cell, but rarely occurred at the top of the cell. These findings could partially explain the absence of CCPs on the top of cells observed in our experiment. We also scanned over the cell top (box height of 15 *µ*m, which is higher than cell height 11 *µ*m averagely) to ensure that this was not caused by incomplete 3D scanning; in some cases, we scanned from top to bottom to exclude the effect of fluorescence quenching. The mechanical and biological mechanisms for this phenomenon need further examination, which are also one of the research focuses of this study. Figure 1 E and F show the scatters of the CCP number and density (number per area) within different size ranges of the 12 cells at the bottom and on the dorsal surface, respectively. Li et al. [12] found that the average CCP density was 2.83*/µ*m^3^. In our research, we determined an average density of 1.54*/µ*m^3^ and 1.02*/µ*m^3^ at the bottom and on the dorsal surface of the total 12 cells, respectively. Although there are more CCPs on the dorsal surface than on the bottom surface, the CCPs on the bottom surface are denser, reflecting that endocytosis processes are more active on the bottom surface than on the dorsal surface. Interestingly, the CCP size and density distribution are heavy-tailed compared with the Normal distribution. A reasonable hypothesis is that the heavy-tailed effect is probably caused by cell membrane tension distribution, which we will also discuss further in this study.

### CCP lifetime observation on cell bottom surface

The lifetime of CCPs observed by TIRF experiments is a well-studied topic with standardized procedures. Our experiment followed the same established protocol as previous studies, detailed in the Methods. Cells were starved before TIRF imaging to ensure an abundance of cargo-free transferrin receptors (TfRs) on the cell membrane. During imaging, transferrins (Tfs) labeled with red fluorescence were added to the cell cultures. Tfs randomly bound to TfRs and were caught by developing CCPs labeled with green fluorescence. This process is illustrated in Figure 2 A. Frames were taken at 2-second intervals, and the video is shown in Supporting Information. Limited by TIRF spatial resolution (*>* 200 nm) [44], only clusters-like of red and green fluorescence can be observed but not the fine size increment of CCPs. The trajectories of CCPs were identified with Imaris software using the same strategies as in the study of Jaqaman et al. [45]. In our cases, a time window of two frames is the optimal gap closing with the frame rate of 0.5 Hz, indicating the end of CCP growth. Since both individual receptors and receptor aggregates generate diffraction-limited image features, the particle positions were detected by local fluorescence intensity maxima and then fitting Gaussian kernels in areas around these local maxima to achieve subpixel localization [42]. When Tf internalized through CCP, the red and green fluorescent trajectories overlapped both temporally and spatially. Thus, CCPs were identified during endocytosis by the temporal and spatial coincidence of the two fluorescent signals. CCPs encountering at least one red trajectory during their lifetime were considered mature, while those without any red trajectory were considered abortive, as illustrated in Figure 2 B. Figure 2 C is a frame from the fluorescence sequence scanned by TIRF (see Video in Supporting Information). Figure 2 C(1) and (2) shows the green fluorescence of CCPs and the red fluorescence of Tfs, respectively; Figure 2 C(3) is the overlapped image of CCPs and Tfs, and Figure 1 C(4) shows the identified trajectories. Loerke et al. [5] studied on cargo and dynamin regulating CCP maturation revealed two short-lived subpopulations corresponding to aborted intermediates and one longer-lived productive subpopulation. They determined that the longer-lived subpopulation was productive because (1) its kinetics matched those of surface-bound transferrin internalization and (2) manual tracking of 450 CCPs yielded a similar average lifetime. Our study focused more on productive CCPs and did not subdivide the abortive groups. We identified 800 trajectories of the productive CCPs and 1012 trajectories of the abortive ones. The CCPs present in the first frame or still existing in the last frame were excluded from the analysis due to incomplete lifetime observation. Figure 2 D and E show examples of typical trajectories of the productive and abortive groups, respectively. The CCPs and the Tfs they interacted with coincided temporally and spatially, as shown in Figure 2 D(1), both exhibiting Brownian-like motion similar to that mentioned by Ehrlich et al. [36]. Figure 2 D(2-3) show the mean square distance (MSD) and diffusion coefficient of CCPs and Tfs. MSD is the average value of the square of the distance that particles or molecules move during a certain period, mathematically expressed as MSD = *<* (*r*(*t*)–*r*(0))^2^ *>*, where *r*(*t*) is the particle’s position at time *t, r*(0) is its position at *t* = 0, and *<>* denotes the average value over time steps. The diffusion coefficient Df and the mean square displacement MSD are related by the Einstein relation, MSD = 4*D*_f_*t* in two dimensions. Figure 2 F (1) shows the spatial distribution of CCP number and lifetime on the cell bottom. Because the lifetime has an exponential distribution, using a linear colormap would concentrate the spot color in the low-lifetime region. Therefore, the colormap is represented by six quantiles of CCP lifetime to better illustrate the relationship between lifetime and CCP number. Figure 2 F (2-3) reveals that CCPs counts and lifetimes were negatively correlated (linear coefficient *r*_P_ = –0.152, *P* = 0.009), consistent with the findings of Willy et al. [21], who noted a shortening of lifetimes with increased initiation and dissolution rates when the cells spread.

**Figure 2:**
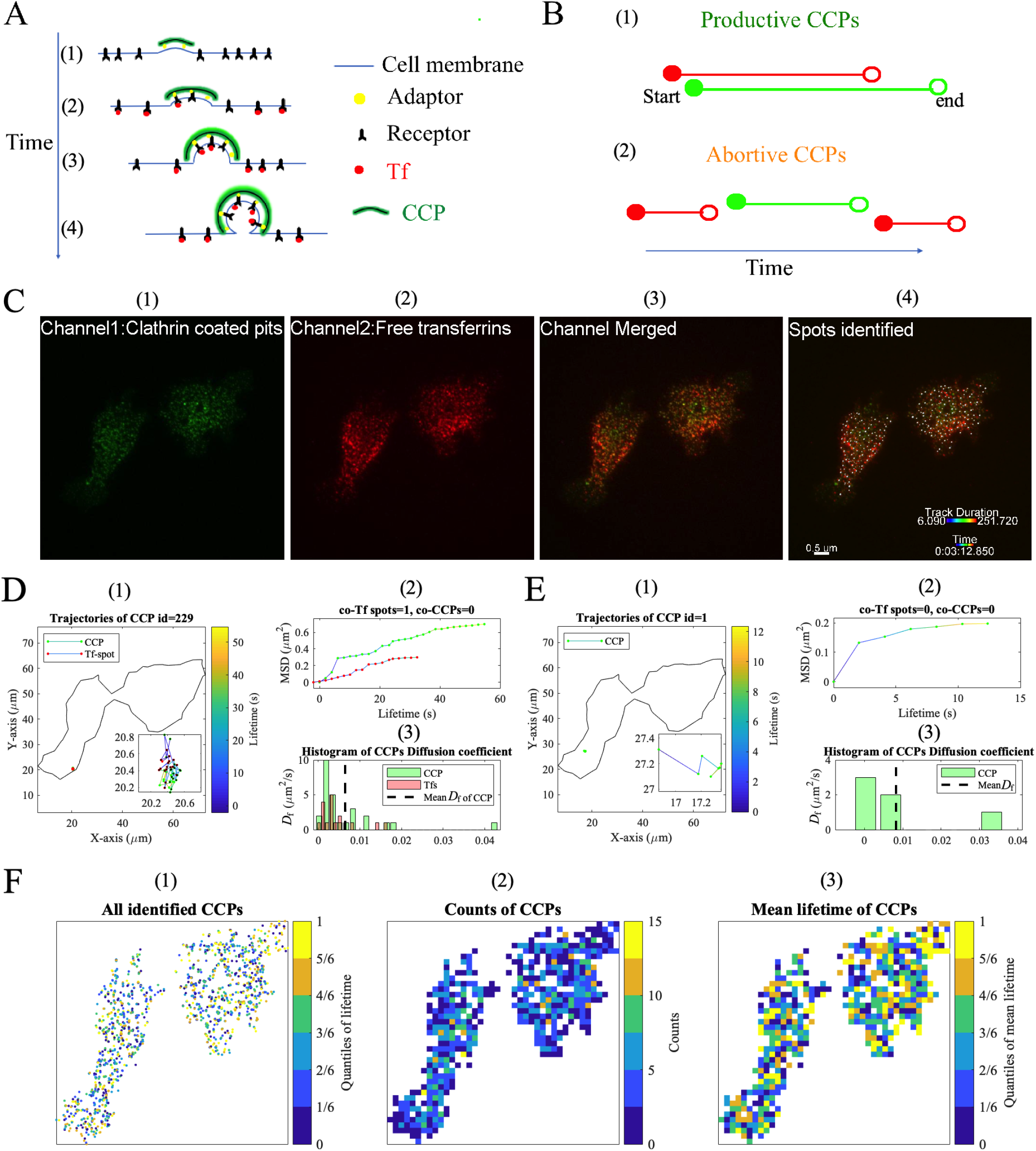
CCP growth time obtained by TIRF. (A) Schematic diagram of CCP wrapping Tfs. (B) CCPs were classified into (1) productive and (2) abortive catalogs. (C) One frame in fluorescent image sequence obtained by TIRF. (1) CCPs. (2)Tfs. (3) Overlapping of CCPs and Tfs. (4) Spots identified by Imaris. Typical trajectories of (E) productive and (F) abortive CCPs. (1) Locations of the trajectories. (2) Mean square distance (MSD) of the trajectories. (3) Histogram of diffusion coefficient *D*_f_. (F) (1) CCPs locations colored by their lifetime, and heatmap of CCP (2) counts and (3) mean lifetime in each mesh grid.

We then analyzed and compared the intensity of CCP generation in the productive and abortive groups to extract key parameters to reproduce the CCP growth process using the Monte Carlo simulation. Figure 3 A shows the new CCPs in each frame. The histogram of the CCP generation intensity is presented in Figure 3B. To determine whether the CCP generation intensity followed Poisson, Rayleigh, or Normal distributions at the 1% significance level, we conducted a Kolmogorov–Smirnov (K–S) test (detailed information is provided in Materials and Methods). The results showed that the Poisson and Rayleigh distributions failed to pass the K–S test, and their fitting curves were not satisfactory, whereas the Normal distributions could did pass the K–S test. The returned value of *H*_0_ = 1 suggested that the K–S test rejected the null hypothesis at the *P <* 0.01 significance level (Table 1). The probability density function (PDF) of CCP generation, fitted by the Normal distribution (Eq.2) with the average and variance values obtained by the maximum likelihood estimates (MLE) are *µ*_c_ = 13.16 *±* 0.80 and *σ*_c_ = 4.78 *±* 0.66 (number of all new CCPs). Figure 3 C shows the discrete and fitted cumulative distribution function (CDF) for a visual comparison of the Poisson, Rayleigh, and normal distributions, which also confirms that the normal distribution is the best. As a result, the normal distribution was used to conduct a Monte Carlo simulation for CCP generation.

**Table 1:**
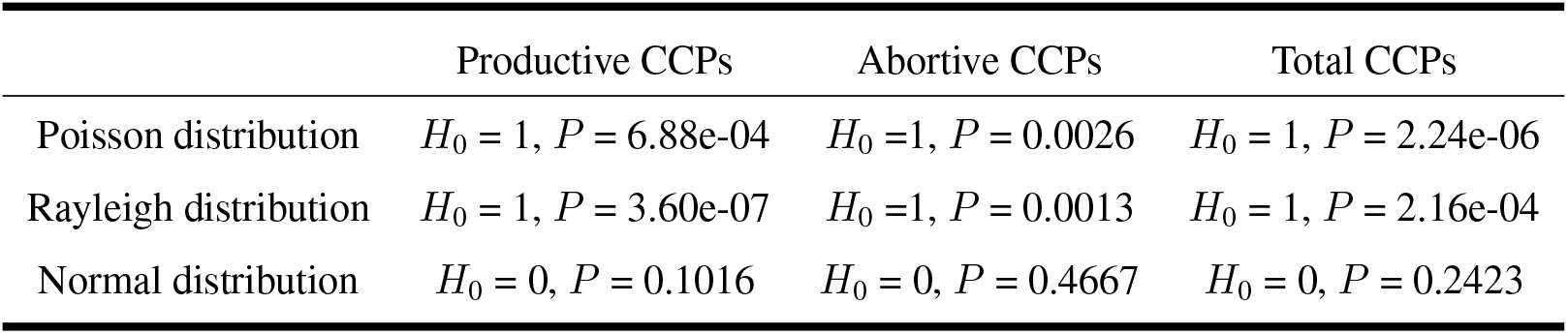
K–S test results of the CCP generation intensity.

|  | Productive CCPs | Abortive CCPs | Total CCPs |
| --- | --- | --- | --- |
| Poisson distribution | $H_0 = 1, P = 6.88e-04$ | $H_0 = 1, P = 0.0026$ | $H_0 = 1, P = 2.24e-06$ |
| Rayleigh distribution | $H_0 = 1, P = 3.60e-07$ | $H_0 = 1, P = 0.0013$ | $H_0 = 1, P = 2.16e-04$ |
| Normal distribution | $H_0 = 0, P = 0.1016$ | $H_0 = 0, P = 0.4667$ | $H_0 = 0, P = 0.2423$ |

**Figure 3:**
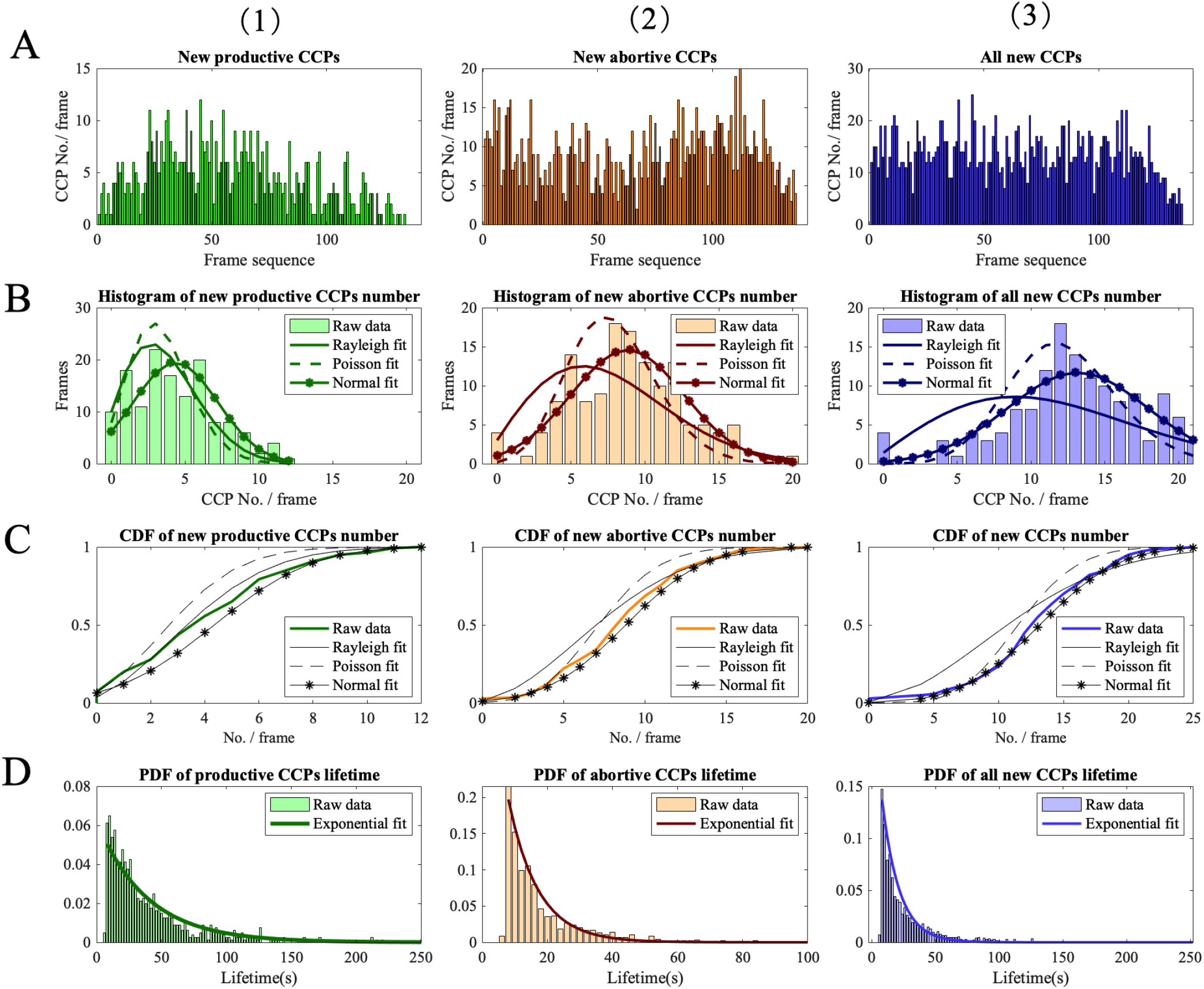
Statistical analysis of CCP lifetime. (A) Numbers of new CCPs in each new frame. (B) Histogram of new CCP numbers in each new frame; they were fitted by the Rayleigh, Poisson, and normal distributions. (C) Cumulative distribution function of the new CCPs in each new frame. (D) Probability density function distribution of the new CCPs in each new frame. (1) Productive CCPs. (2) Abortive CCPs. (3) All CCPs.

Figure 3 D shows the PDF of the new CCPs in each new frame. The exponential distribution (Eq. 9) was used to estimate the expectation of CCPs’ lifetimes as shown in Figure 3 A(2-3). The expected lifetime of productive and abortive CCPs, maximum likelihood estimates, were 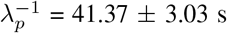 and 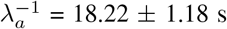, respectively. The longest observed lifetime for productive and abortive CCPs were 250 s and 100 s, respectively, reflectingsignificant individual variability in CCP lifetime. Loerke et al. [5] classify CCPs by piecewise fitting the joint PDF of the CCP lifetimes to categorize CCPs into three groups: productive, early abortive, and late abortive CCPs. The abortive group was divided into early abort (5.2 *±* 0.1 s) and late abort (15.9 *±* 1 s), respectively. The lifetime of the productive CCPs was 86.9 *±* 5.8 s, which followed an exponentially decaying distribution. The differences in cell lines used in their study (monkey kidney epithelial cells, BSC1) versus our study (RBL-2H3 cells) might explain the slight differences in average CCP lifetimes. Interestingly, Yang et al. [46] classified CCPs into those exhibiting clathrin waves and those without. They reported that the lifetime of CCPs that exhibited clathrin waves (median: 37.4 *±* 11.1 s; mean:32 *±* 42 s) was consistently shorter than that of CCPs without clathrin waves (median: 72.5 *±* 18.3 s; mean: 48 *±* 38 s) in RBL-2H3 cells. As far as know, Dynamin is often employed to verify the completion of endocytosis, which is a GTPase integral to the final stage of clathrin-mediated endocytosis [28]. Grassart et al. [47] demonstrated that dynamin recruitment occurs in two distinct phases: an initial recruitment phase followed by a burst of dynamin activity just before scission. This observation aligns with Loerke et al.’s [5] identification of different stages of CCP maturation, including productive and abortive pathways. Consequently, even though this study did not utilize dynamin to mark the final stage of endocytosis, the analysis of cargo and CCP dynamics and lifetimes remains sufficient to identify the productive group.

### Relationship between the CCP size and lifetime

We reproduced the growth process of CCPs by Monte Carlo simulation, with the PDFs summarized from our experimental data (Algorithm 1). The proportion of the abortive CCPs was determined to be *γ* = 0.56, based on the ratio of 1012 abortive CCPs to a total of 1812 CCPs identified. Results from 10,000 times Monte Carlo simulations are presented as mean and standard deviation, shown as error bars in Figure 4 A. These simulation results closely match the experimental results, validating the assumptions that the generation intensity of CCPs follows a normal distribution and their lifetime distribution follows an exponential distribution. The proportions of CCPs in different size range are displayed in bar charts in Figure 4 B. The average CCP size proportions are similar at the bottom and on the dorsal surface (Figure 1 E and F), although slight variations exist for individual cells (see Supporting Information). CCPs with size between 100–250 nm (represented by cyan, green, and yellow) account for 62.4 % of the total CCP population, which is the majority subpopulation of the total CCPs. CCPs smaller than 100 nm account for 12.0 %, while those larger than 250 nm constitute 25.6 %. In our Monte Carlo simulation, we investigated two potential relationships between CCP size and growth time (time from formation to current growing size), which were *t* ∝ *R*^2^ and *t* ∝ *R*^3^ base on mathematical intuition, where *R* is the radius of CCP. These relationships were considered because CCPs are usually considered as shell structures mechanically [48] and they are fruit-like in shape. The average and variance distributions of simulation results for *R*^2^ and *t* ∝ *R*^3^ are represented by red and green error bars, respectively. Conversely, the green error bars, representing the *t* ∝ *R*^2^ relationship, closely match experimental data, peaking between 150 and 200 nm. This indicates that CCP growth time is likely proportional to their volume. Given the SIM resolution of approximately 40–60 nm, and considering that a fluorescent spot must consist of at least four pixels to be recognized as a CCP, distinguishing CCPs smaller than 100 nm from noise is challenging. This systematic error may explain the substantial discrepancy between experimental data and Monte Carlo simulations for sizes smaller than 100 nm.

**Figure 4:**
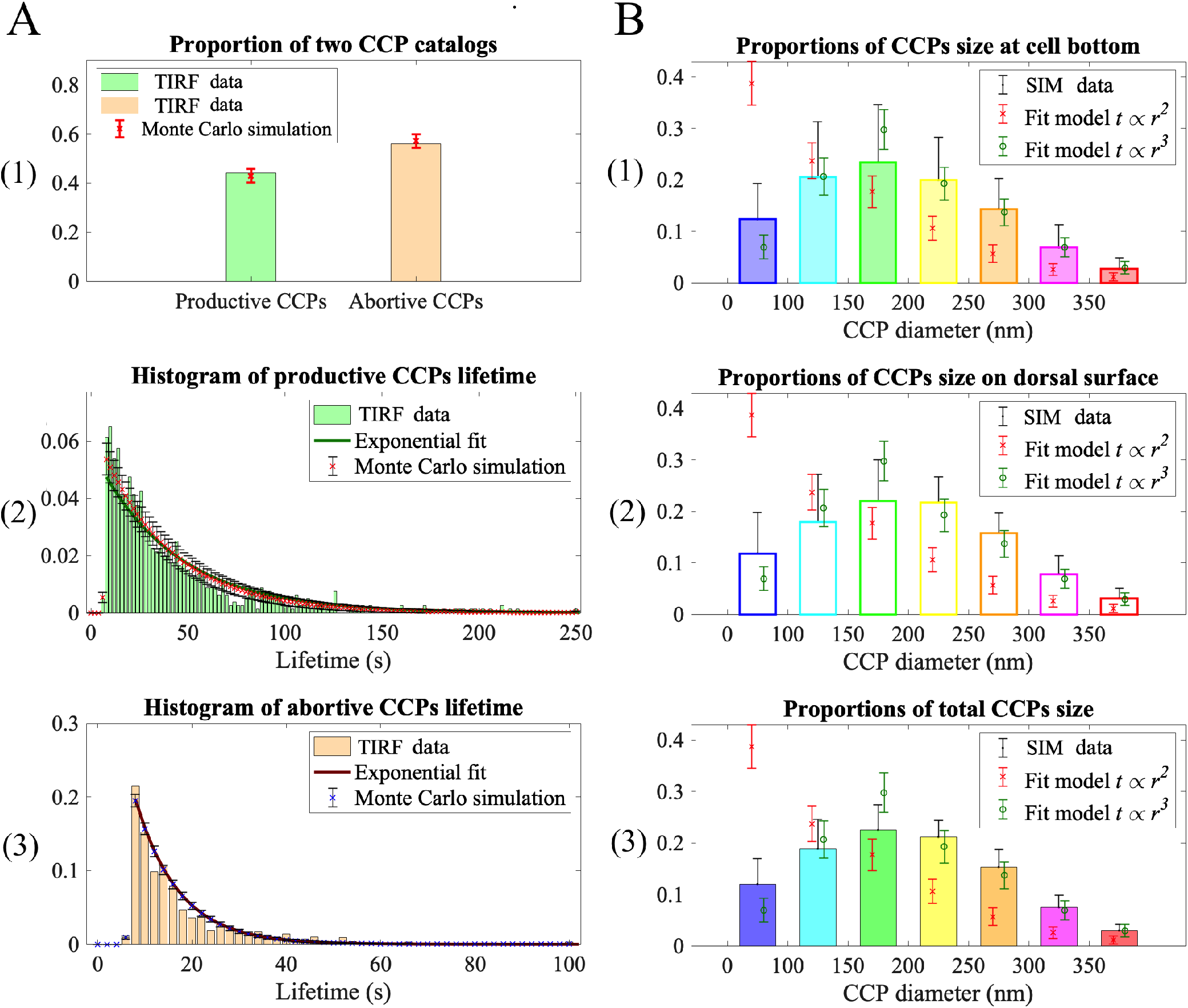
CCP growth process is reproduced by Monte Carlo simulation and compared with the TIRF and SIM experimental information. (A) Proportions of the productive and abortive CCPs and their lifetime histogram. (B) Comparison of the CCP proportions in different size ranges, considering two possible relationships between the CCP size and growth time.

Figure 5 shows the analyzed results of the relationship between the CCP size and lifetime. We terminated the Monte Carlo simulation at *t* = 600 s, which was the total observation time of the TIRF image sequence and sufficiently long. Figure 5 A displays the histogram of the growth time of the CCPs still developing when the Monte Carlo simulation terminated. In Figure 2 and Figure 4 we excluded the CCPs still existing in the last frame due to incomplete lifetime observation; however, here we counted them as their growth time since they were still developing. This comparison with the CCP growth time observed by TIRF further confirmed that the Monte Carlo simulation scheme was reliable and that the relationship *t* ∝ *R*^3^ was reasonable. Figure 5 B shows the fitting curve between the CCP size and growth time, where the growth times corresponding to the selected sizes are labeled on the axis and highlighted in colored dash lines. Among these, the CCP growth time with a size *<* 100 nm was only 4 s, indicating that the CCPs were captured soon after they were initiated, consistent with the actual situation. Figure 5 C shows the CDF of the CCP growth time by Monte Carlo simulation, revealing that CCPs with a size *<* 200 nm had a corresponding growth time of *>* 24 s, which accounted for 60%. The proportion of CCPs with a size *>* 250 nm and a corresponding growth time *>* 76 s was less than 23%, which was also consistent with the experimental observation of 25.6% for the CCPs size *>* 250 nm. Figure 5 D presents the CDF of the growth time of productive and abortive CCPs, with the corresponding PDF is already shown in Figure 4 A(2–3). In Figure 5 D, the lower abscissa is labeled with the growth time, and the upper abscissa is labeled with the corresponding CCP sizes as simulation using the relationship *t* ∝ *R*^3^. It shows that 95% of the abortive CCPs were of a growth time within 32 s, and only 0.2% of abortive CCPs were of size *>* 250 nm. Loerke et al. [5]also presented a subset of long-lived, isolated CCPs, which were highly likely (99%) to represent productive events. The predicted maximum size of abortive CCPs in our study is within 200–250 nm, compared to 90 nm predicted by Banerjee et al. [8], due to the observation size effect of a fluorescent microscope as mentioned earlier. Therefore, when investigating the relationship between the cell membrane tension distribution and CCP size, this 0.2% of abortive CCPs could be neglected, and the CCPs with a size *>* 250 nm were considered productive. Overall, we categorized CCPs into three groups based on the relationship between the CCP size and growth time: (1) size *<* 100 nm were de novo CCPs mixed with some image noises, (2) size between 100–250 nm were developing CCPs, including partial abortive ones, and (3) size *>* 250 nm were mature CCPs in the productive group.

**Figure 5:**
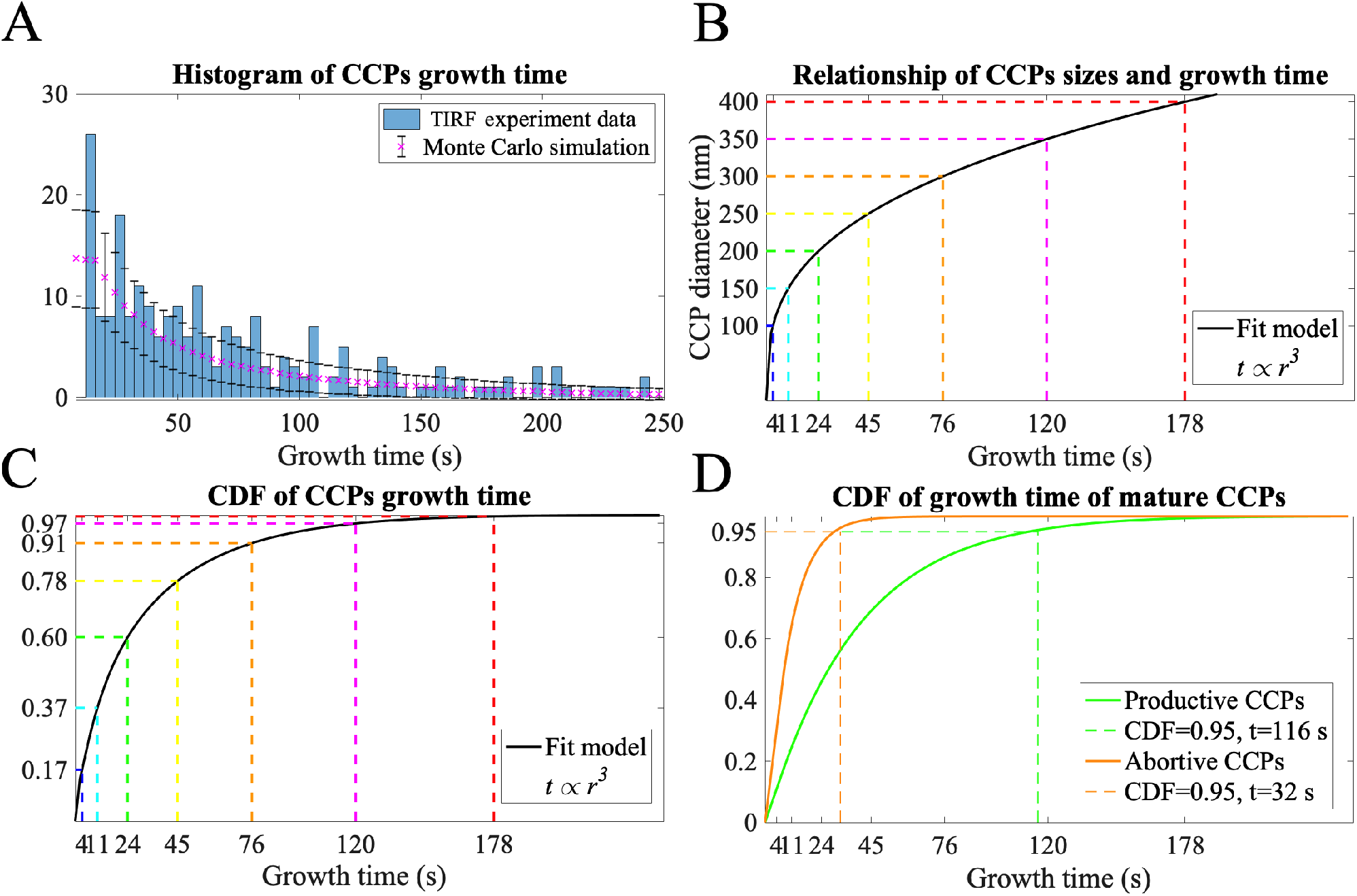
Growth time and sizes of the developing CCPs when the Monte Carlo simulation terminated. (A) Histogram of the CCP growth time compared with the experimental values. (B) Relationship between the CCP size and growth time fitted by *t* ∝ *R*^2^. (C) CDF of the growth time of the total simulated CCPs fitted by *t* ∝ *R*^3^. (D) CDF of the CCP growth time of experimental data with double abscissas fitted by *t* ∝ *R*^3^.

### Apparent Membrane tension distribution

The apparent membrane tension of a cell corresponds to the combination of in-plane and cortical tension [49, 50]. The lipidbilayer beers in-plane tension due to the osmotic pressure difference across the plasma membrane, while the cytoskeleton bears the cortex tension. The apparent membrane tension, which includes both contributions, results from the multitude of binding interactions between cytoskeletal elements and membrane phospholipids [17, 51]. The apparent tension is mainly determined by cell deformation, whereas in-plane tension receives the influence of CCP growth. Thus, the apparent tension serves as the background tension for CCP growth. In this study, for simplicity, the term “membrane tension” is used to refer to the apparent membrane tension, as the lipid bilayer detachment is not considered. Directly measuring the spatial distribution of membrane tension on cells via existing experimental methods remains a challenge. In our previous study [38], cell spreading was assumed to be a quasi-static process, and the dynamic process of cell spreading from the suspension state was solved inversely through an algorithm that alternately executed a particle-based method and searched for the minimum deformation energy. A standard spread cell model was established by 3D reconstruction of the cell shapes, as shown in Figure 6 A(1), representing the average geometry of 12 scanned cells (see Figure 1). The cell-spread shapes were obtained until the cell deformation energy function converged to the minimum, and the membrane tension distribution was predicted using Hooke’s law, with the membrane was discretized into triangular grids. The first principle tension describes the maximum tensile force of the triangular element in mechanics. Tan et al. [19] confined the cells growing on micropatterns through fibronectin, presenting a quantitative analysis of how physical cues, specifically cell spreading area, altered the dynamics of CCPs. Their results showed a positive correlation between the average cell membrane tension and the spread area. Cells with larger spreading area have more short-lived CCPs but a higher CCP initiation rate. Inspired by their study, we calculated a square cell with the same bottom as the standard spread cell model and a circular cell with twice bottom area, as shown in Figure 6 A(2-3). Pietuch and Janshoff [52] utilized atomic force microscopy to obtain the tension changes in cell membranes during spreading. They found that the cortical tension of the cell membrane first increases from a low value and, after 30 min of cell spreading, the apparent tension is stabilized at a mean value of (0.6 *±* 0.05) mN/m. Qualitative and quantitative comparisons with previous experimental results in references [19,52] validated the reliability of our numerical method as referenced in [38]. Our simulation results demonstrated that the membrane tension gradient demonstrated decreases from the bottom to the top of the cell. Cells with larger bottom areas developed greater tension compared with cells with smaller bottom areas, and the cells with the same bottom area but a larger perimeter also experienced higher membrane tension. Furthermore, this numerical method can be applied to estimate the membrane tension distribution of cells in any free-spreading shape (see Figure 6 A(4-6) and Supporting Information). The tension is higher where the filamentous pseudopodia grow. Several reports indicated that the membrane tension mechanically resisted shape changes involving an increase in the total surface area of the cell. For example, the expansion of cellular protrusions was resisted by membrane tension during cell spreading [17, 53] and cell migration [54, 55]. Therefore, our estimates of tension distribution were consistent with these findings.

**Figure 6:**
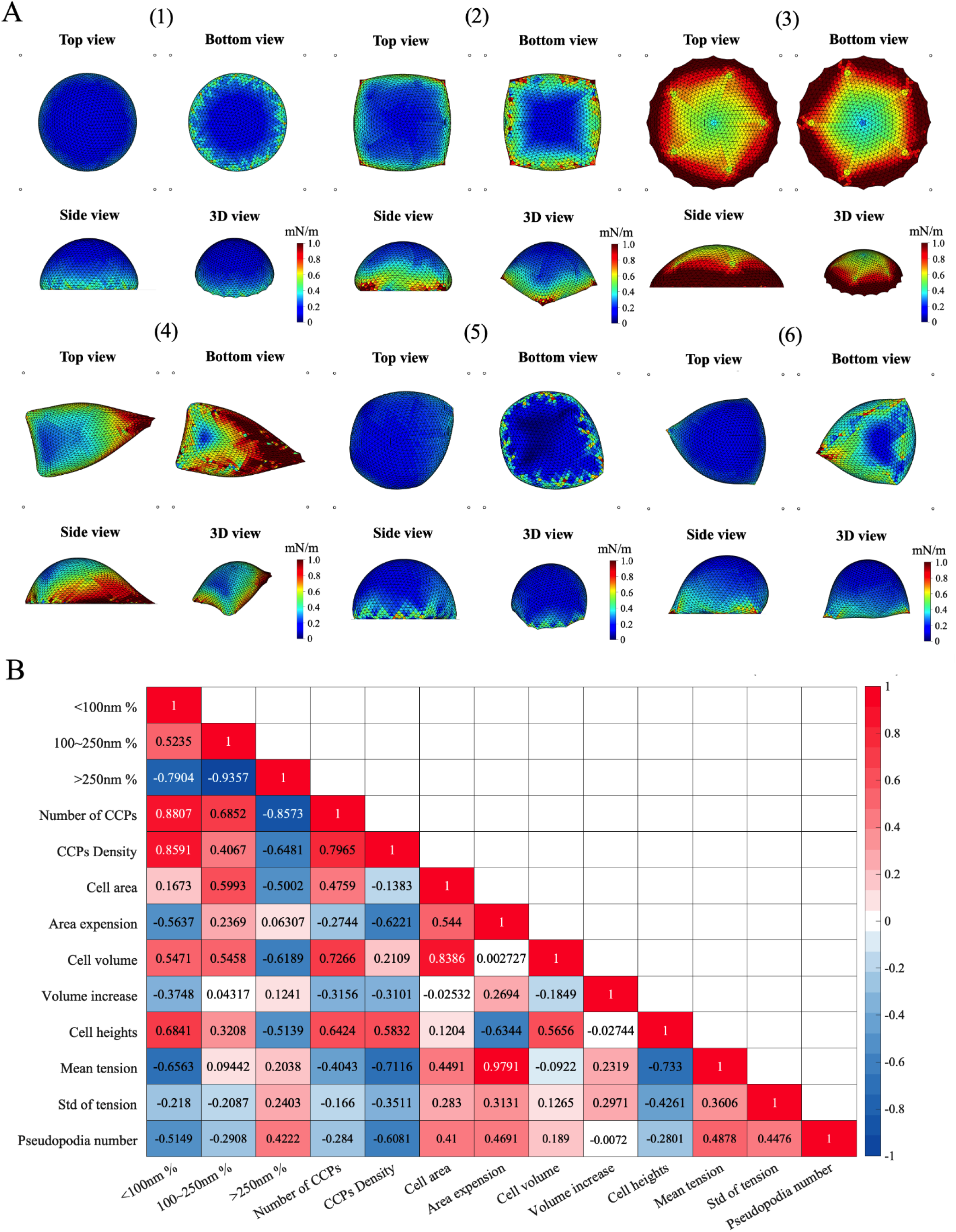
Apparent Membrane tension distribution and the relevance analysis of CCP information. (A) Numerical results of the first principal membrane tension of (1) the standard spread cell model, (2) a square-bottomed cell with the same bottom area, and (3) a circular cell with a double-bottomed area compared with the standard spread cell model. (4–6) Cell ID.01-03. Cell ID.04-12 can be seen in Supporting Information. (B) Relevance of the CCP number, cell geometry, and membrane tension properties.

We examined the relevance of CCP number, geometric parameters, and tension across these 12 cells, and calculated the correlation matrix using the Pearson correlation coefficient *r*_P_ (Eq. 10) as shown in Figure 6 B. The corresponding values of these parameters are listed in Table S1 in Supporting Information. The parameters characterizing the quantity of CCPs included the CCP number, percentage of different size ranges, and density. The parameters characterizing the geometric properties of the cells included area, volume, and height. The parameters characterizing cell membrane tension included the average and variance of tension, area expansion, volume increase, and number of pseudopodia. Interestingly, the number of pseudopodia was positively correlated with the average tension because the cells needed to consume more energy for each new pseudopodium they extended. Groups of parameters representing similar properties show the same positive or negative trends in correlation, including “the number and the density of CCPs” as the quantity information, “the cell area, the cell volume, and the cell height” as the geometry information, “the area expansion and the volume increase of cells” as the deformation information and “the average and the variance of tension” as the tension information. Results show that the total CCP number was positively correlated with the geometric properties such as the cell area and volume, but negatively correlated with the membrane tension, consistenting with the findings of Dai et al. [15]. In other words, the larger-sized cells had a higher CCP number, and the cells in a higher-tension state possessed a lower CCP number. It is worth mentioning that the membrane tension is positively correlated with the mature CCP percentage, and this is a new finding in our study. We will explain this interesting discovery in the following section.

### Relationship between the spatial distribution of CCPs and membrane tension

In this section, we primarily discuss the patterns of CCP size and the spatial distribution of cell membrane tension. Cell ID.11, whose bottom surface was nearly circular, was selected to compare the CCPs locations and the membrane tension distribution, as its tension distribution was close to that of the standard spreading cell model, as shown in Figure 7 A(1–3). Data for ther cells are provided in Supporting Information. The CCP size histogram displayed a heavy tail compared with the normal distribution [Figure 7 A(4)]. The local membrane tensions were sorted into six quantiles (Figure 7 B). The histogram of the membrane tension was nonuniform, with concentrations between 1/6 and 3/6 of the maximum tension [Figure 7 B(4)], because membrane tension is non-linear distributed. Addtionally, CCPs of different sizes were all concentrated within a range of 0.15–0.35 mN/m of background membrane tension [Figure 7 C(1-2)]. Therefore, we sorted the CCPs by six quantiles of the membrane tension [Figure 7 C(3–4)]. On the bottom surface, fewer CCPs were found in 0–1/2 quantiles of tension than in 1/2–1 quantiles of tension. On the dorsal surface, CCPs with a size *<* 250 nm exhibited a relatively uniform distribution, while CCPs with a size *>* 250 nm appeared concentrated at higher-tension quantiles.

**Figure 7:**
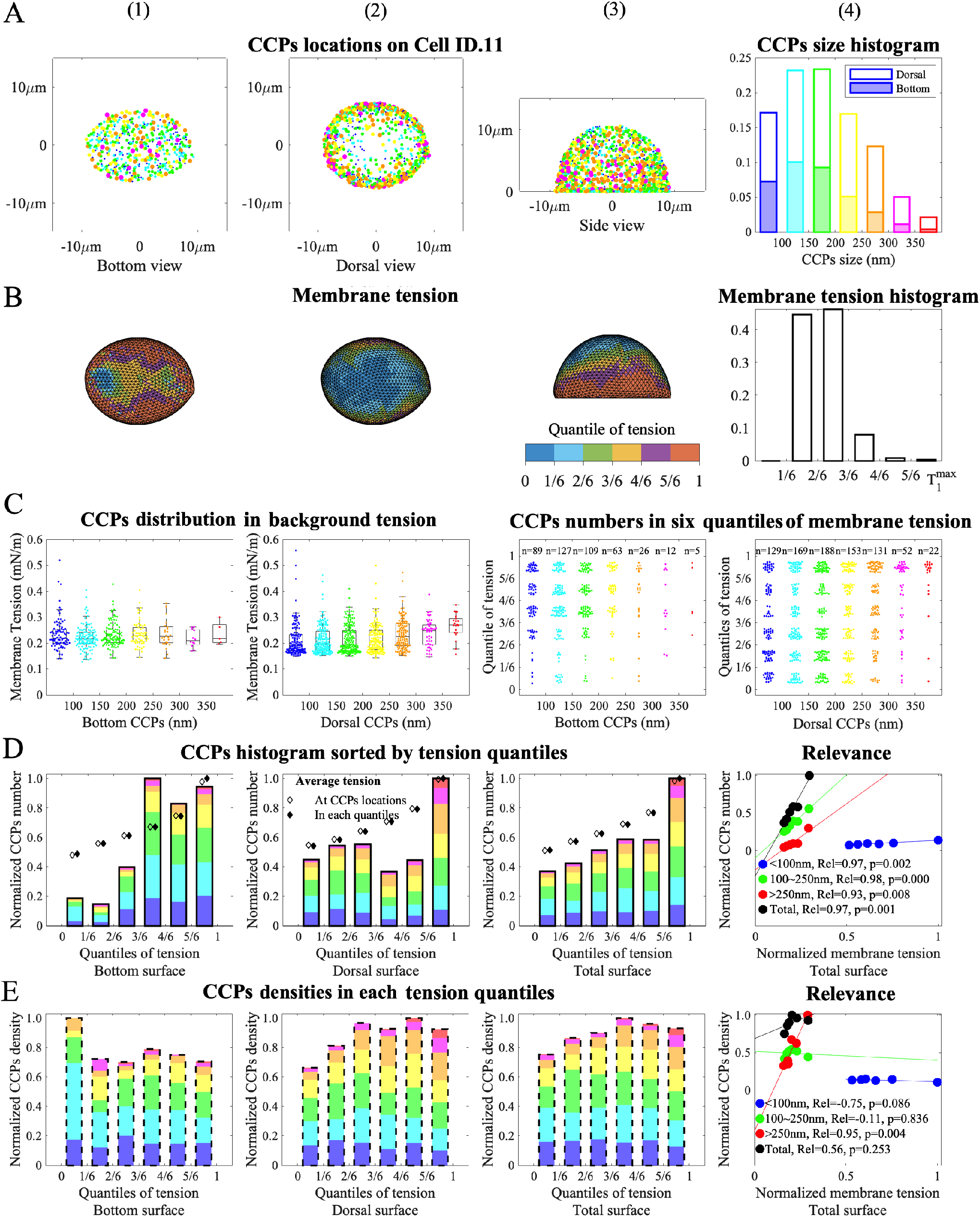
Comparison of the CCP locations and the membrane tension on Cell ID.11. (A1–3) CCP locations on the surface scanned by SIM. (A4) Stacked histogram of the CCP sizes on the dorsal and bottom surfaces. (B1–3) Simulated membrane tension distribution in six quantiles. (B4) Membrane tension histogram. (C1–2) Background membrane tension distribution where CCPs are located and cataloged by their size ranges. (C3–4) CCP numbers in six quantiles of the membrane tension cataloged by their size ranges. (D1–3) A stacked histogram of the CCP sizes was sorted by six tension quantiles. Spots ♦ and ♢ represent the average of the area expansion rate in each quantile and within the local areas of the CCPs, respectively. (D4) Relevance of the CCP numbers in each quantile and the average tension where CCPs were located. (E1–3) CCP density bar in each tension quantile. (E4) Relevance of the CCP density in each quantile and the average tension where CCPs were located.

We further compared the CCP number bar across six quantiles of membrane tension. The CCP numbers were normalized by their maximum values at the bottom [Figure 7 D(1)], on the dorsal surface [Figure 7 D(2)]), and the total surface [Figure 7 D(3)]. We used two methods to normalize the average area expansion rate (physically representing the tension): by the background tension where the CCPs are located, and by the tension intervals where the CCPs are located, represented by “spots ♦” and “spots ♢,” respectively. The CCPs number stacked bars show a similar increasing trend with both spots ♦ and ♢, and they are better fitted to the histogram of CCPs on the total surface [Figure 7 D(3)] than that at the bottom [Figure 7 D(1)] or on the dorsal surface [Figure 7 D(2)]. Moreover, spots ♦ and ♢ are almost overlapped, indicating no obvious tension selectivity for the CCP locations. The other cells show similar trends as the Cell.ID 11, as presented in Supporting Information (Figure S1-11), indicating that the appearance of larger quantities of CCPs in the higher-tension region is likely a general pattern, especially a sharp increase in the CCP numbers in 5/6–1 quantile of tension, a reason for which is discussed later.

As analyzed earlier, the CCPs can be divided into de novo, developing, and mature categories according to the relationship between their size and growth time. Thus, we further analyzed the correlation between the CCP number and the area expansion rate, as shown in Figure 7 D(4). The blue, green, and red dots represent the de novo, developing, and mature CCPs, respectively. The *R*_el_ and *P* -value are the coefficient of determination (Eq. 11) and significance, respectively. The total CCP number on the surface was positively correlated with the average membrane tension where the CCPs were located. The CCPs size ranges of 100–150 nm and 150–250 also show a positively correlated with the membrane tension. However, CCPs smaller than 100 nm exhibit variability across different cells.As analyzed in Figure 4 and Figure 5, We found that shorter-life CCPs were of smaller sizes. Besides, the low signal-to-noise ratio at the top of the cell due to light scattering, and therefore the error in the data analysis, was higher in this CCP category. More importantly, a lower expected number of CCPs appearing in the lower-tension region, as shown in Figure 7 D(1-3), the CCP density distributions were checked to confirm whether it was due to a smaller area of this tension region. If so, the density distributions should be close to the uniform distribution. But no specific pattern of CCPs density distributions with tension was observed in six quantiles, as shown in Figure 7 E(1–3), and the correlation analysis supported this [Figure 7 E(4)]. The other cells also had a random distribution of CCP densities and no significant statistical correlation between densities and tensions, indicating that the initial location of CCPs are independent of membrane tension, but their final growth sizes are related to tension. (see Supporting Information).

We found an interesting phenomenon: a sharp increase in CCP numbers in 5/6–1 quantile of tension. To determine whether this was specific to the Cell.ID 11 or a general rule, we examined the Pearson correlation coefficient between the CCP number and the average tension for each cell, as shown in Figure 8. Although the CCP number did not always monotonically increase within the 0–5/6 quantile of tension, a sharp increase was observed in both the CCP number and the average tension within the 5/6–1 quantile for all cells (Figure 8 A, B and c). Additionally, there was a slight variation in the percentage of different CCP sizes, with a slight decrease in the percentage of CCP *<* 250 nm and a slight increase in the percentage of *>* 250 nm (Figure 8D), again suggesting that more mature CCPs appeared in the higher-tension region. The CCP number and the percentage of CCPs *>* 100 nm showed significant correlations with the average tension across six quantiles for all the cells, but the densities of CCPs did not (Figure 8 E and F).

**Figure 8:**
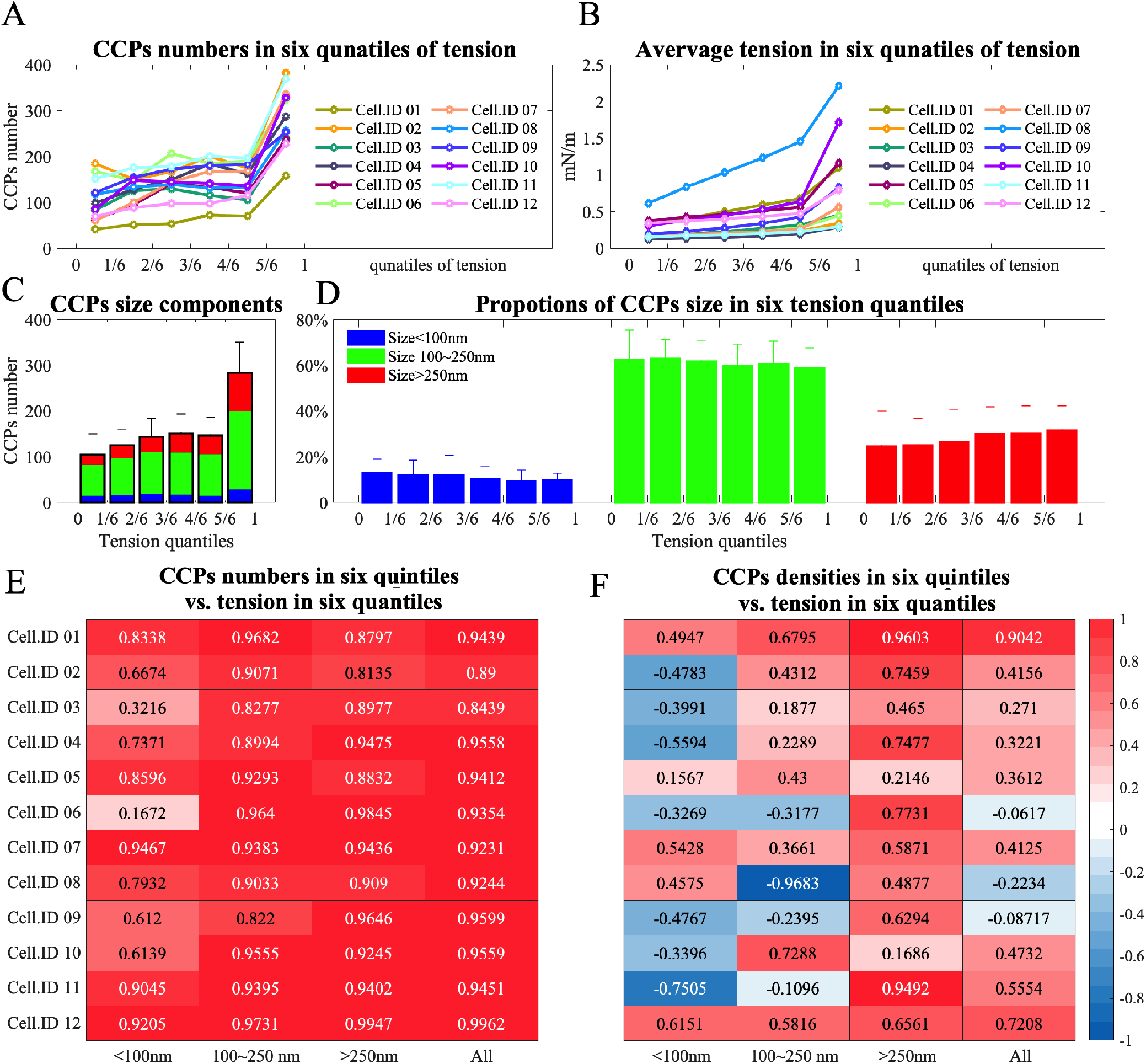
The correlation between the CCP number and the average tension for each cell in our experiment and numerical results. (A) The CCP numbers and (B) the average tension in six quantiles of tension. (C) The component of the CCP size and (D) the comparison of their proportion changes in six quantiles of tension. (E) The relevance of the CCP number and (F) the density to the average of tension in six quantiles.

### Effect of apparent membrane tension on CCP mature size

To analyze how cell membrane tension affects CPP mature size, we previously modeled the CCP as a shell structure with bending rigidity [39]. An optimization algorithm was proposed for minimizing the total energy of the system, which included the deforming nanoparticle (red), the receptor–ligand bonds (black), the cell membrane (blue), and the CCP structure (green), as shown in Figure 9. The CCP structure enables full wrapping of the nanoparticles at various membrane tensions represented by the pre-elongation rate *I*_e_, which is a parameter to describe the degree of background tension. *D* represents the size of the nanoparticle, and the minimum searching algorithm needs an guessed initial CCP size *D*_c0_, that can precisely encapsulate the nanoparticle to start the prediction [Figure 9 A-D(1)]. Figure 9 E shows the normalized sizes of mature CCPs (*D*_c_*/D*_c0_) increased in a similar trend with *I*_e_ increase, no matter what *D*_c_ was. When *I*_e_ *>* 1.2, most of the receptor–ligand bonds in the wrapping area broke because their lengths exist the broken length as CCP size increase. As the membrane tension increased, the normalized deformation energy also increased in similar trends (Figure 9 F), regardless of the nanoparticle size. Therefore, the CCP mature size was dependent on the membrane tension rather than on the size of the nanoparticle. Since more energy was needed for CCP maturation under higher membrane tension, it was easy to understand that the total CCP number decreased as the average tension increased (Figure 6). However, under high-tension conditions, larger CCPs are required to ensure successful endocytosis and facilitate the dynamin-assisted “necking” stage.

**Figure 9:**
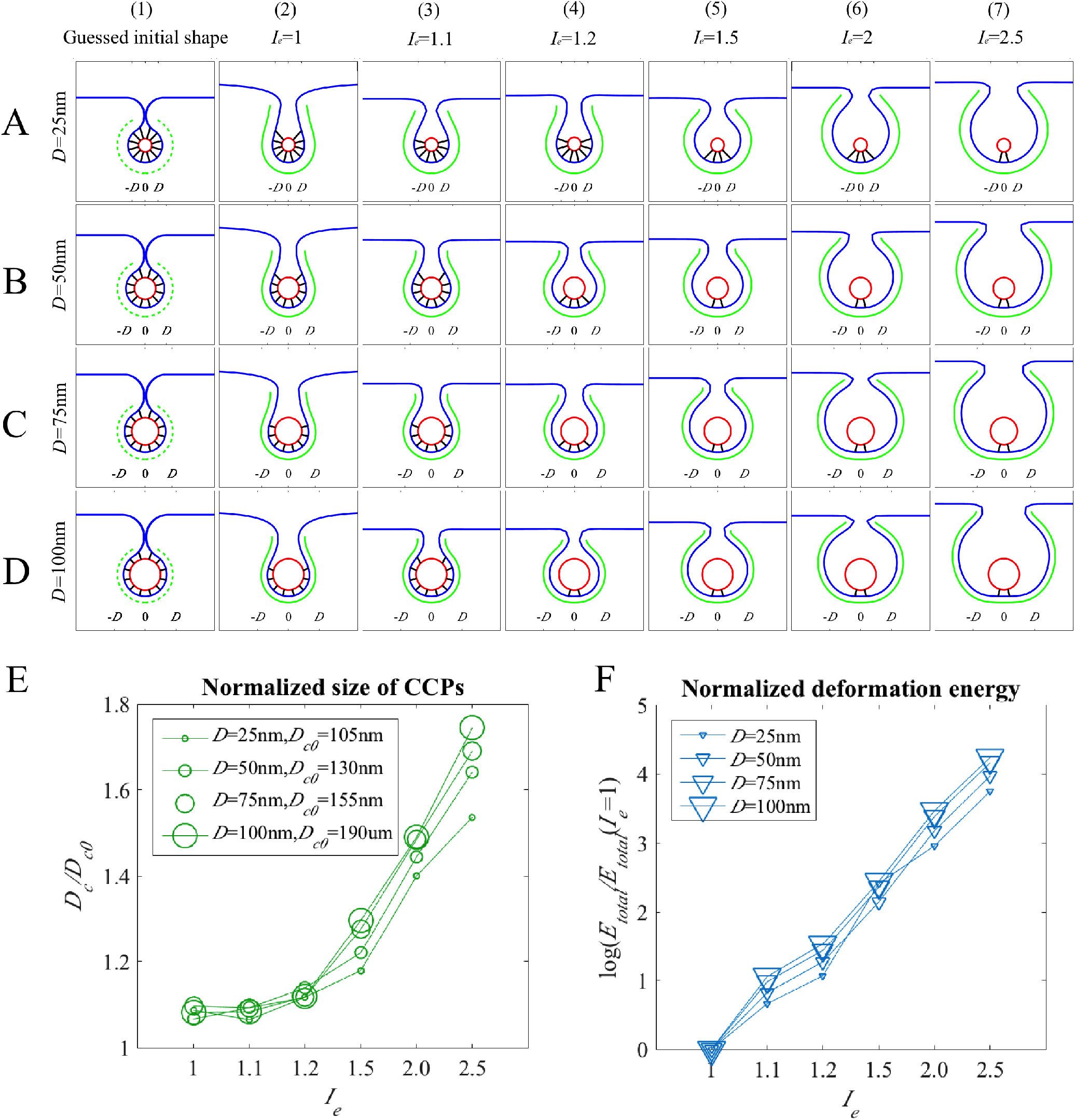
Size of mature CCPs increase with cell membrane tension when wrapping nanoparticles of different diameters. (A) D = 25 nm, (B) D = 50 nm, (C) D = 75 nm, and (D) D = 100 nm. (E) CCP size increased with the membrane tension. (E) Deformation energy increased with the membrane tension.

## Discussion

### Why we use SIM

Huang et al. [56] utilized the stochastic optical reconstruction microscopy (STORM) to reveal the three-dimensional CCPs, demonstrating their half-spherical cage-like morphology at the nanoscale level. Aguet et al. [57] used the lattice light-sheet microscopy (LLSM) to study the dynamic behavior of CCPs and changes in cell morphology during cell division and found that CCPs form at a lower rate during late mitosis. In their study, the scanning precision in the XY-axis and Z-axis were about 100 nm and 250 nm, respectively, which makes it challenging to obtain accurate measurements of CCP size.

STORM achieves extraordinary spatial resolution, around 20-30 nm laterally and 50-60 nm axially. This enables precise reconstruction of nanoscopic structures without sample scanning. However, STORM’s temporal resolution is a significant drawback, often requiring minutes to hours for a complete image, making it less suitable for studying dynamic processes in living cells. LLSM excels in rapid, high-resolution 3D imaging with minimal photodamage, capturing fast dynamics in living cells with excellent temporal resolution. However, its spatial resolution, while good axially (around 250 nm), does not match the lateral resolution of STORM or SIM. Additionally, LLSM requires complex and computationally intensive deconvolution.

Given our study’s requirements for both high spatial and rapid temporal resolution to accurately capture the CCPs size of a whole cell, we opted for SIM. In this study, it achieves 60 nm lateral and approximately 250 nm axial resolution by illuminating the sample with patterned light and reconstructing high-resolution images. Its relatively high speed, about 30 minutes to scan a whole cell, makes it ideal for capturing CCPs size across multiple cell samples in a single experiment. Therefore, SIM is a necessary trade-off between temporal and spatial resolutions. Nevertheless, the results of SIM are still countable and reasonable for CCP size variation analysis.

### Energy considerations in Membrane tension and CCPs curvature

The diversity of membrane curvature modes in CCP formation and maturation is a significant area of research. It has already long been suggested that high membrane tension leads to excess exocytosis, an increase in cell surface area, and a decrease in tension, and vice versa [15]. The mechanisms behind CCP curvature changes exhibit some contradictory behaviors, particularly in how they are influenced by membrane tension and energy dynamics. High membrane tension can inhibit the transition from flat to curved structures, acting as an energetic barrier that stabilizes flat clathrin lattices. This was highlighted by Bucher et al. [58], who noted that high tension increases the lifetimes of clathrin structures and prevents their curvature changes. Similarly, Boulant et al. [2] demonstrated that increased membrane tension, whether through hypo-osmotic shock or mechanical stretching, impedes CCP curvature changes. Conversely, when membrane tension is reduced, CCPs can more readily transition to a curved state. This dual behavior showcases how membrane tension can both stabilize flat structures and, when decreased, facilitate the formation of curved clathrin pits [59].

From an energy viewpoint, the bending of clathrin lattices involves overcoming the energetic constraints imposed by the membrane and the clathrin coat itself. Sochacki et al. [60] pointed out that the concentration distribution of different proteins in the edge and interior regions of the CCPs changes with the formation and maturation of the CCPs, and this change is consistent with the trend of clathrin changes and is related to the surface area of the CCPs. Therefore, cell membrane tension and energy may affect the morphology and function of CCPs by regulating the distribution of different proteins in the interior and edge of the CCPs. Various proteins contribute to membrane bending by decreasing membrane rigidity or by acting through steric bulk to induce curvature. ENTH/ANTH and BAR domain-containing proteins, such as epsins and CALM, play roles in this process by reducing membrane rigidity and assisting in the transition from flat to curved structures [27, 61, 62].

The relationship between membrane tension and energy dynamics is complex and interdependent. High membrane tension poses an energetic barrier, requiring additional forces or modifications (like actin recruitment) to achieve membrane curvature. On the other hand, energy dynamics involving protein interactions and membrane properties dictate how efficiently these transitions occur under varying tension conditions. However, why the CCPs concentrate in the highest-tension range of each cell, especially the mature CCPs (essentially the molecular mechanism of how clathrins aggregate more in a higher-tension region), is still unclear. The concentration of mature CCPs in high-tension regions likely results from the competition and interplay of several factors, including membrane tension, actin dynamics, the recruitment of specific adaptor and curvature- sensing proteins, membrane composition, and energetic considerations. These factors collectively create an environment that supports the aggregation and maturation of clathrin-coated pits.

### Future Research Prospects

To further investigate the mechanism by which CCPs concentrate in high-tension regions, especially in mature stages, several experimental and modeling approaches can be pursued. Here are some suggestions and future research prospects: Utilize super-resolution microscopy techniques, such as STORM and photoactivated localization microscopy (PALM) to observe the dynamic changes of CCPs on the cell membrane. These techniques provide sub-cellular level resolution, allowing detailed analysis of CCP behavior in different tension regions. Employ genetically encoded membrane tension sensors (such as FLIPPER-TR or Piezo1) to measure changes in cell membrane tension in real-time, because these sensors can provide spatiotemporal distribution information of membrane tension and correlate it with CCP distribution.

Improving the spatial resolution of LLSM can significantly benefit the study of CCPs. Advances such as adaptive optics to correct for sample-induced aberrations, or incorporating techniques like super-resolution microscopy into light-sheet setups, can provide clearer, more detailed images of CCP dynamics in three dimensions. This would allow for more precise tracking of CCP formation and maturation under varying membrane tensions.

Integrate multiscale models, from molecular-level dynamic simulations to cell-level mechanical models, to construct a comprehensive model of CCP formation and tension regulation mechanisms. By combining experimental data and simulation results at different scales, a more holistic understanding of CCP mechanisms can be achieved. Focus on studying the dynamic changes during CCP formation and maturation, especially the spatiotemporal distribution and behavior under high- tension conditions. Develop real-time imaging and dynamic simulation methods to capture CCP behavior in various tension environments.

## Conclusions

This study utilized SIM to obtain the size and spatial distribution of CCPs in fixed RBL-2H3 cells, and TIRF microscopy to observe CCP lifetimes in living cells. The generation intensity of CCPs fits a normal distribution, as confirmed by the K-S test. Monte Carlo simulations accurately reproduced the CCP formation and maturation/abortion processes, with CCP lifetimes following an exponential distribution. Results indicate that CCP growth time is proportional to their volume, and CCPs can be categorized into de novo, developing, and mature stages based on size. There is a significant positive correlation between membrane tension and the size of mature CCPs, particularly in the highest tension regions. This correlation supports the numerical prediction that CCPs grow larger to overcome higher energy barriers caused by increased membrane tension. Additionally, CCP numbers are positively correlated with cell area and negatively correlated with membrane tension, indicating that higher membrane tension hinders CCP maturation. These findings provide valuable insights into the role of membrane tension in CCP maturation and dynamics.

## Acknowledgments

This study was supported in part by the National Natural Science Foundation of China (12072198, 12202258) and the China Scholarship Council for Joint PhD program (201206230004). Wang Xi received funding from the Mechanobiology Institute seed grant, Ministry of Education’s Academic Research Fund Tier 1 Grant (R-397-000-247-112). RBL-2H3 cells stably expressing clathrin light chain A tagged with green fluorescent proteins were provided by Professor Min Wu’s research group at the National University of Singapore. The experimental work was conducted at the Nanolab of Professor Chwee Teck Lim’s research group at the National University of Singapore.

## Materials and Methods

### Cell culture

RBL-2H3 cells stably expressing clathrin light chain A tagged with green fluorescent proteins were cultured in minimum essential medium (MEM) (Invitrogen 11095-09) with 20% fetal bovine serum (FBS) (Sigma f4135-500 mL) and 50 *µ*g/mL gentamicin (Invitrogen 15750-060). The cell culture conditions were 5% carbon dioxide, 37^°^C, and 95% humidity.

### Fixed-cell 3D scanning using SIM

After culturing in a normal medium for 24 h, the cells were washed three times with phosphate-buffered saline (PBS) (pH = 7.4) and fixed with 4% paraformaldehyde (pH = 7.4). SIM (n-sim; Nikon) is with objective lens CFI apoTIRF 100*×* oil (N.A. 1.49), 488-nm laser transmitter, and CCD camera (DU-897, Andor and Technology). The horizontal (XY-axis) and the vertical (Z-axis) scanning step sizes were 24 nm and 120 nm, respectively. Imaris software was used to identify green fluorescent spots and obtain their size and XYZ coordinates. It was considered that the fluorescent spots on the surface of the cell were the CCPs undergoing endocytosis, whose position was extracted using the self-built MATLAB code.

### CCP interaction with Tf observed using TIRF

After culturing in a normal medium for 24 h, the cells were cultured in starvation solution without FBS for 4 h and then transferred to the image buffer for time series imaging to eliminate the influence of endocytosis signals from other proteins. During imaging, 20 *µ*g/*µ*L Tf Alexa Fluor 647 (Thermo Fisher) solution was added. Fluorescent images with two channels were taken every 2 s. Trajectories of red and green spots were identified using Imaris software. The temporal and spatial co-location analysis of the trajectories of red and green fluorescence was programmed using the self-built MATLAB code.

### Statistical analysis

K-S test is a non-parametric test used to determine whether a sample comes from a specific probability distribution, in this case, the normal distribution. The test statistic *D*_ks_ is calculated as the maximum absolute difference between the empirical cumulative distribution function (ECDF) of the sample data and CDF of the theoretical normal distribution. The formula to compute the test statistic *D*_ks_ for a one-sample K-S test for normality is:

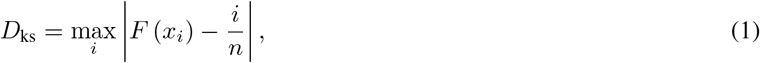

where *F* (*x*_*i*_) is the CDF of the theoretical normal distribution evaluated at the *i*-th ordered observation, 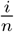 is the proportion of observations up to the *i*-th ordered observation, *n* is the sample size, and max_*i*_ denotes the maximum absolute difference taken over all ordered observations.

The significance of the *D*_KS_, statistic is evaluated against critical values from the Kolmogorov distribution to determine whether the observed data significantly deviates from the normal distribution. If the calculated p-value *P* = 1 − CDF(*D*_KS_) is greater than a predetermined significance level (e.g., *P* = 0.05), the null hypothesis of normality is not rejected (*H*_0_ = 0), indicating that the observed data can be reasonably modeled by a normal distribution. Conversely, a *P* -value below the threshold indicates rejection of the null hypothesis, suggesting that the observed data significantly differ from a normal distribution.

To estimate the parameters of a normal distribution using MLE, we start with the PDF of the normal distribution, which is given by:

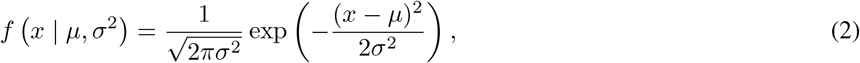

where *x* is the observed data, *µ* is the mean (location parameter), and *σ*^2^ is the variance (scale parameter).

The likelihood function *L* (*µ, σ*^2^ | *x*) for a set of independent and identically distributed (i.i.d.) observations *x*_1_, *x*_2_, …, *x*_*n*_ from a normal distribution is the product of the PDF evaluated at each observation:

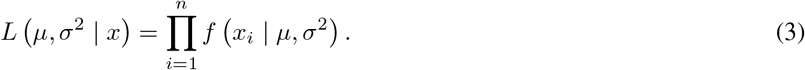

Taking the natural logarithm of the likelihood function (log-likelihood) simplifies the computation:

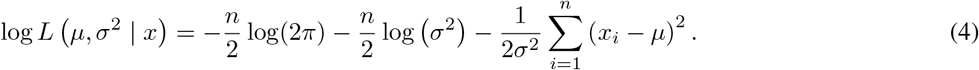

To find the maximum likelihood estimates 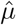 and 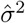, we differentiate the log-likelihood function with respect to *µ* and *σ*^2^, respectively, set the derivatives equal to zero, and solve for the parameters:

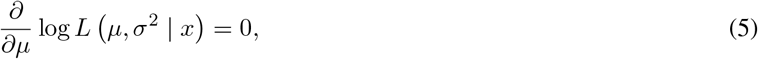

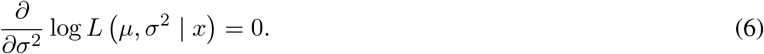

Once the derivatives are equated to zero, the resulting equations can be solved numerically to obtain the maximum likelihood estimates of *µ* and *σ*^2^, These estimates represent the parameter values that maximize the likelihood of observing the given data under the assumed normal distribution model.

Similarly, MLE can be used to estimate the parameters of the Poisson *P* (*X* = *k*), Rayleigh *f* (*x*; *σ*), and exponential *f* (*x*; *λ*) distributions:

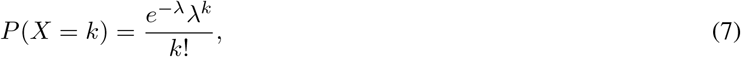

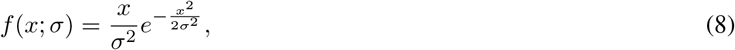

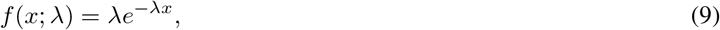

where *λ* is the average rate parameter, and *σ* is the scale parameter.

The Pearson correlation coefficient measures the linear relationship between two continuous variables. It ranges from -1 to 1, where *r*_P_ = 1 indicates a perfect positive linear relationship, *r*_P_ = 1 indicates a perfect negative linear relationship, *r*_P_ = 0 indicates no linear relationship. The formula to compute the Pearson correlation coefficient *r*_P_ is

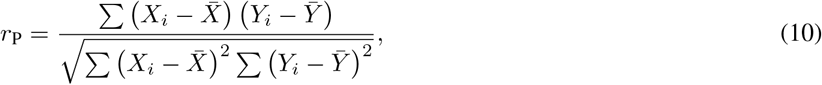

where *X*_*i*_ and *Y*_*i*_ are individual data points for variables *X* and *Y*, 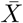 and 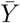 are the means of variables *X* and *Y*, respectively.

Arrange the computed correlation coefficients into a symmetric matrix, known as the correlation matrix. The diagonal elements of the matrix will be 1, as each variable is perfectly correlated with itself. The off-diagonal elements will represent the correlations between pairs of variables.

The correlation coefficient *r*_P_ will indicate the strength and direction of the linear relationship between the variables. Use linear regression analysis to model the relationship between the independent variable *X* and the dependent variable *Y*. Fit a linear equation of the form *Y* = *β*_0_ + *β*_1_*X* + *ϵ*, where *β*_0_ is the intercept (y-intercept), *β*_1_ is the slope (regression coefficient), *ϵ* is the error term.

Assess the goodness of fit of the linear regression model using metrics such as the coefficient of determination *R*_el_:

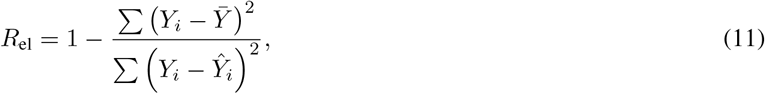

where 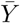 is the mean of the observed values of the dependent variable and *Ŷ*_*i*_ is the predicted value of the dependent variable for the *i*-th observation.

*R*_el_ quantifies the proportion of the variance in the dependent variable that is predictable from the independent variable. A value of *R*_el_ close to 1 indicates a strong linear relationship between the variables.

The *P* -value for the linear correlation between two sets of data *X* and *Y* can be calculated using the Pearson correlation coefficient. The calculation involves transforming the correlation coefficient into a *t*-statistic form, and the *P* -value can then be obtained using the cumulative density function of the *t*-distribution. Let *n* be the sample size The formula for the *t*-statistic is:

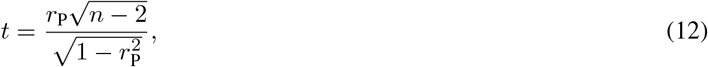

where *n* is the sample size. Subsequently, the *P* -value can be obtained using the CDF of the *t*-distribution with the appropriate degrees of freedom (*df* = *n* - 2).

#### Algorithm 1

Monte Carlo simulation of CCP generation

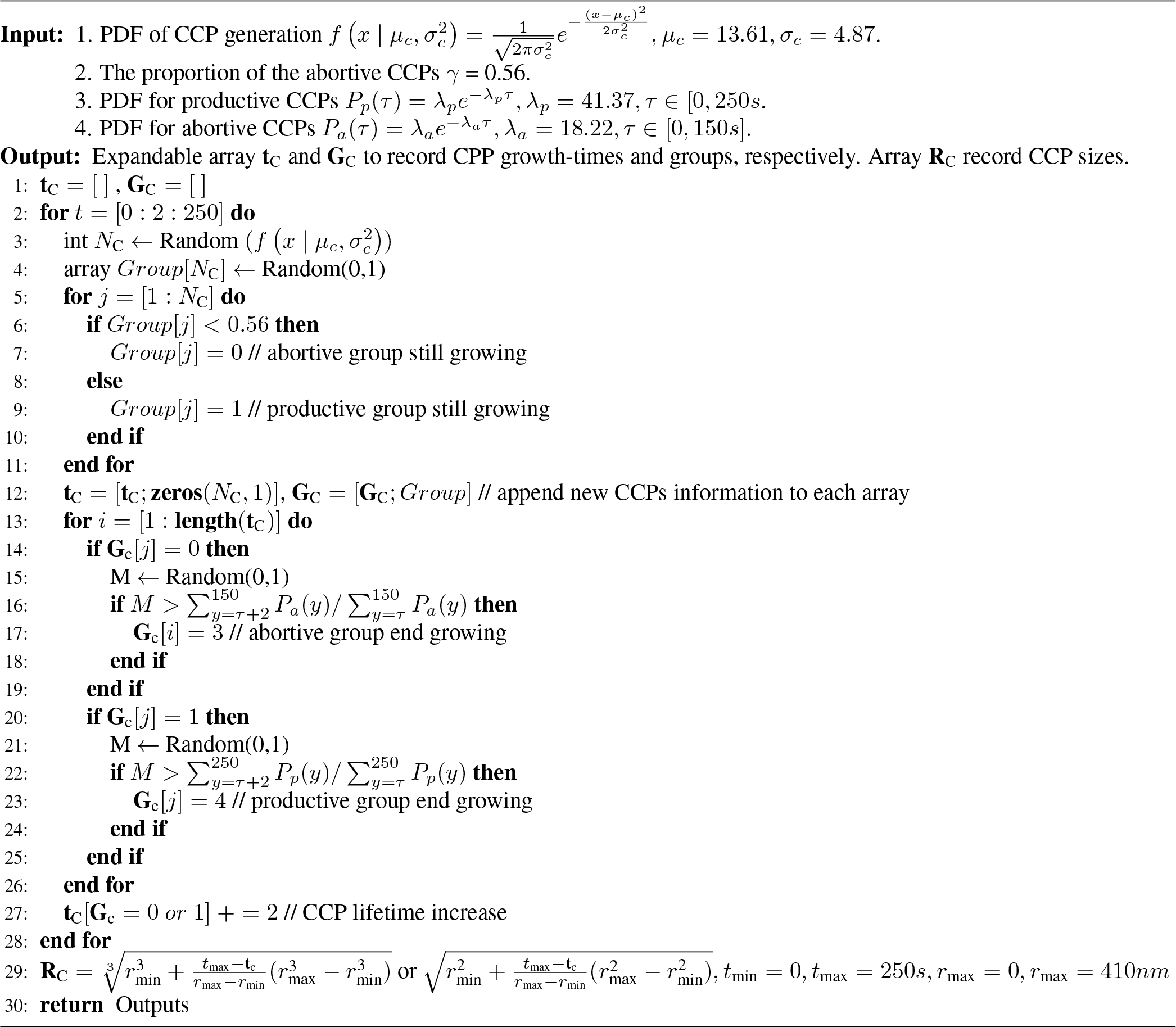

## Abbreviations

CCP: Clathrin-coated pit
CDF: Cumulative distribution function
K-S: test Kolmogorov–Smirnov test
LLSM: Lattice light-sheet microscopy
MLE: Maximum likelihood estimation
MSD: Mean square distance
PDF: Probability density function
PSF: Point spread function
SIM: Structured illumination microscopy
STORM: Stochastic optical reconstruction microscopy
Tf: Transferrin
TfR: Transferrin receptor
TIRF: Total internal reflection fluorescence microscopy

## Supporting Information

### Rowdata:https://data.mendeley.com/v1/datasets/pfgfbv3rjx/draft?preview=1

**Figure S1.**
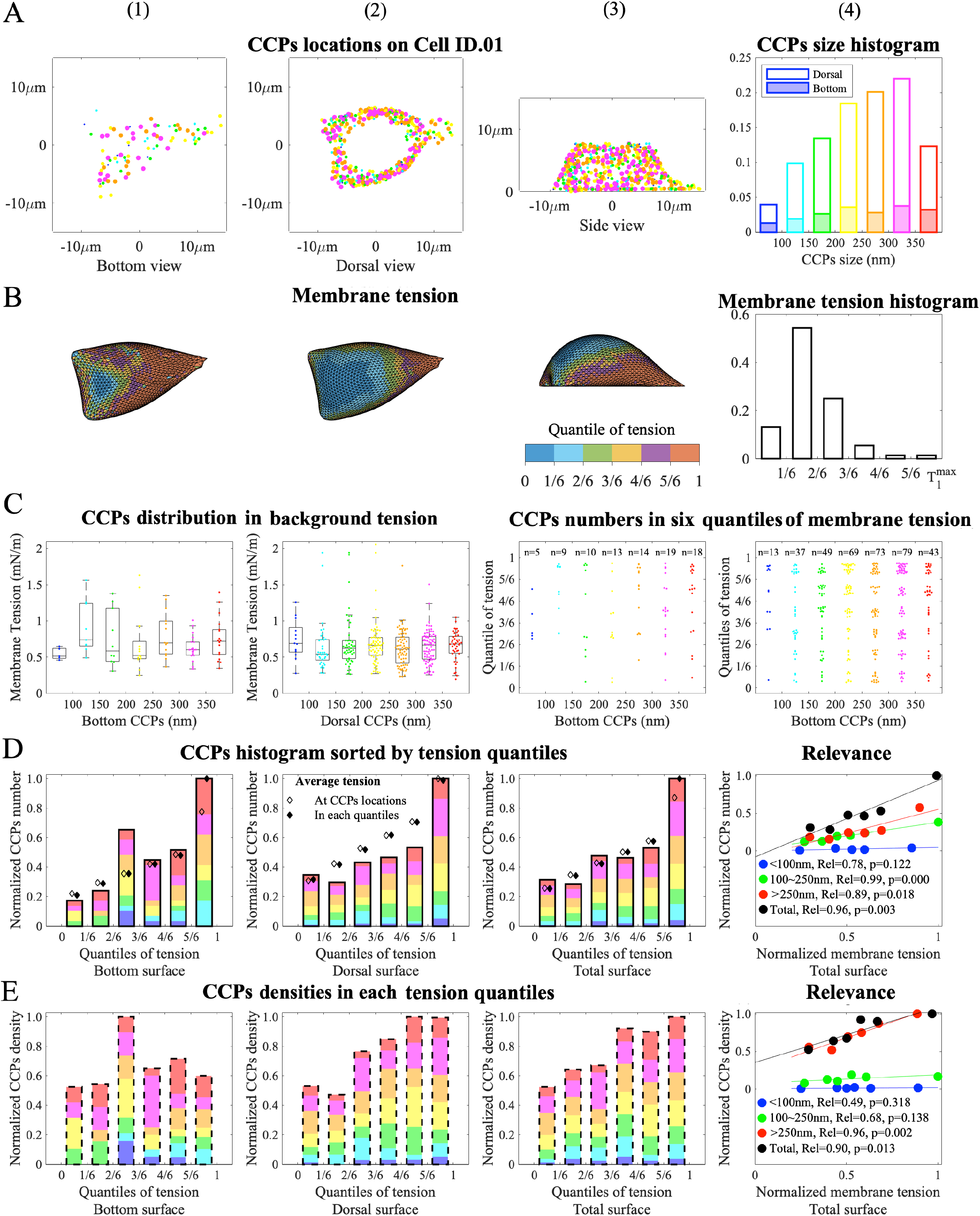
The comparison of the CCPs locations and the membrane tension on Cell ID.01.

**Figure S2.**
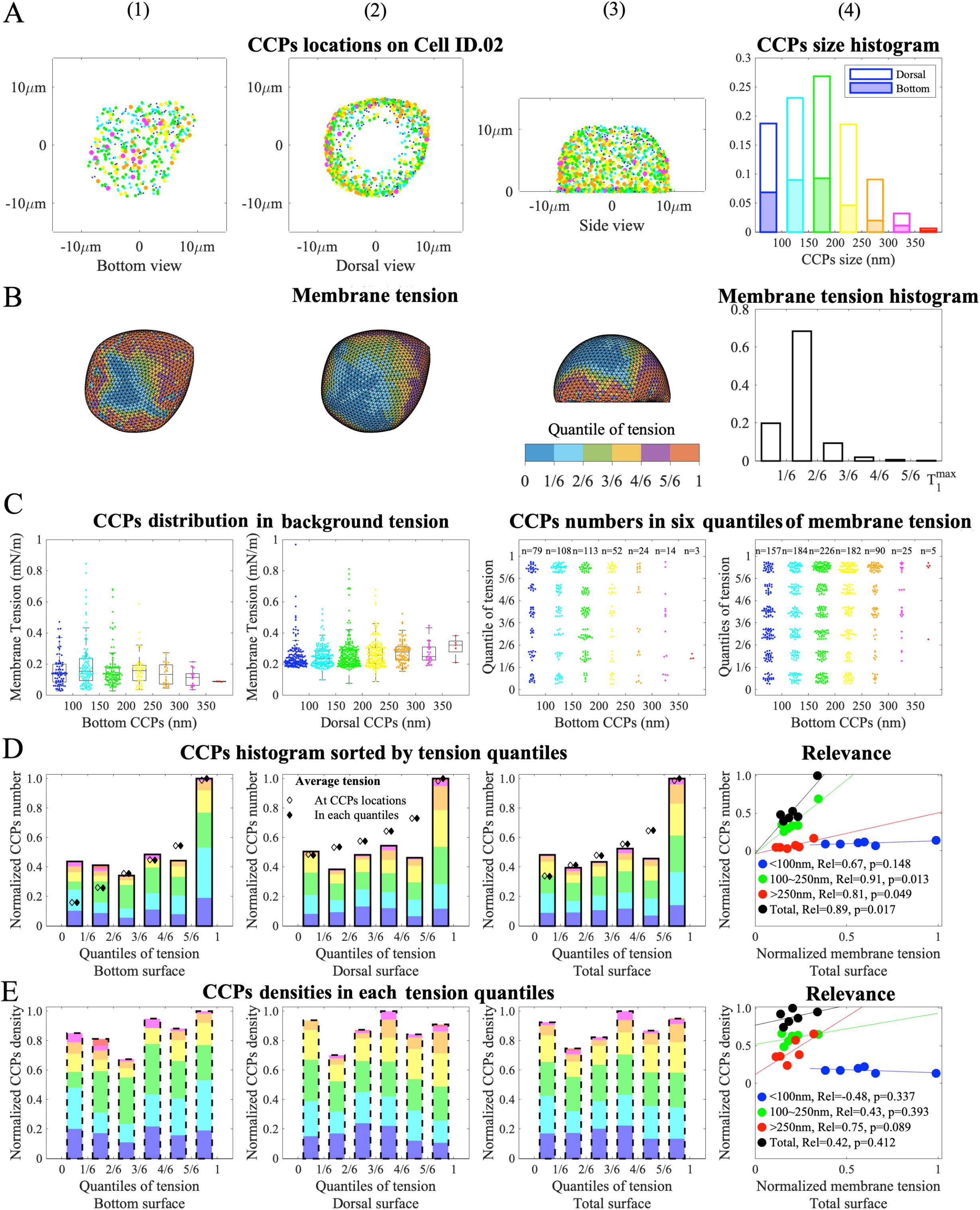
The comparison of the CCPs locations and the membrane tension on Cell ID.02.

**Figure S3.**
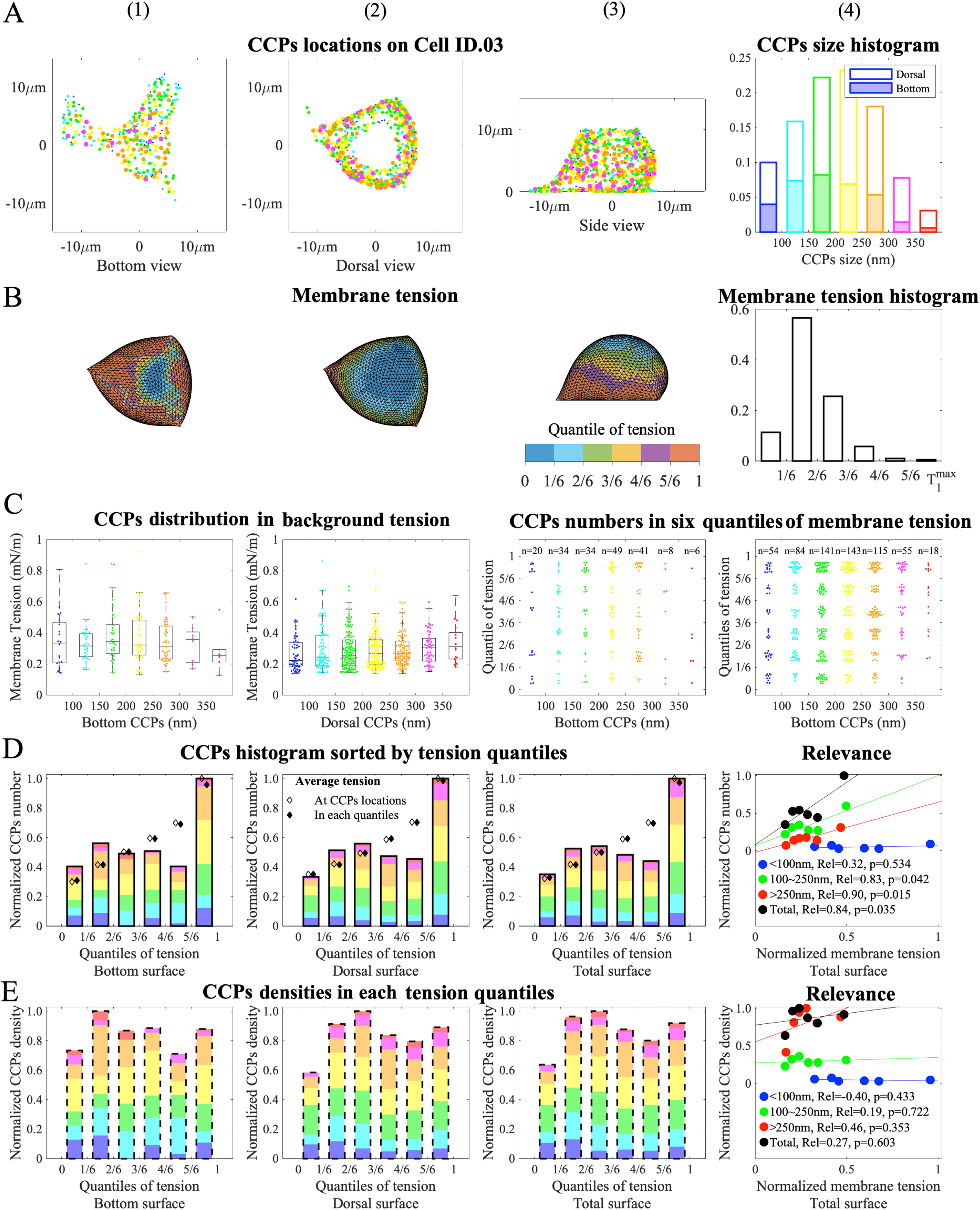
The comparison of the CCPs locations and the membrane tension on Cell ID.03.

**Figure S4.**
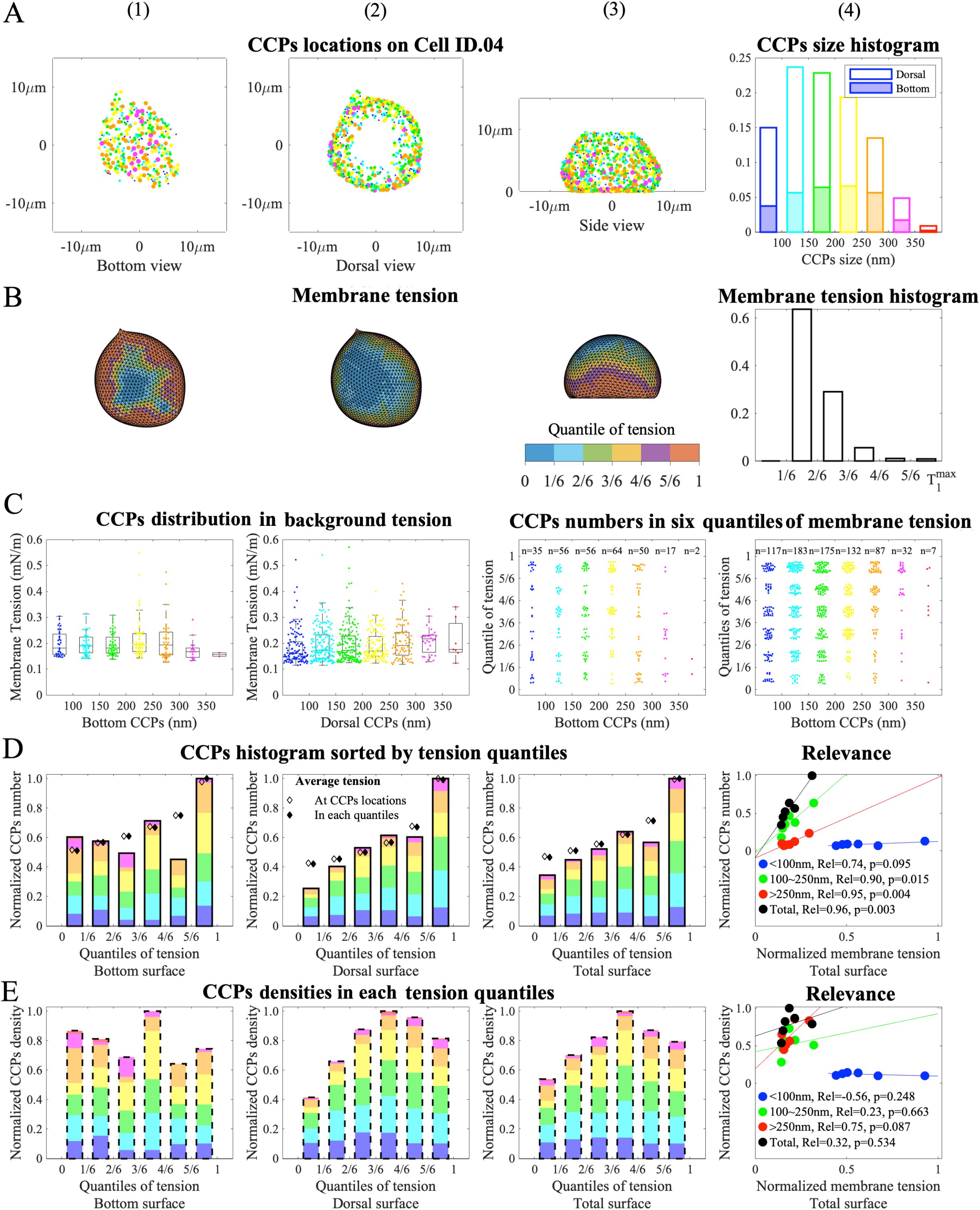
The comparison of the CCPs locations and the membrane tension on Cell ID.04.

**Figure S5.**
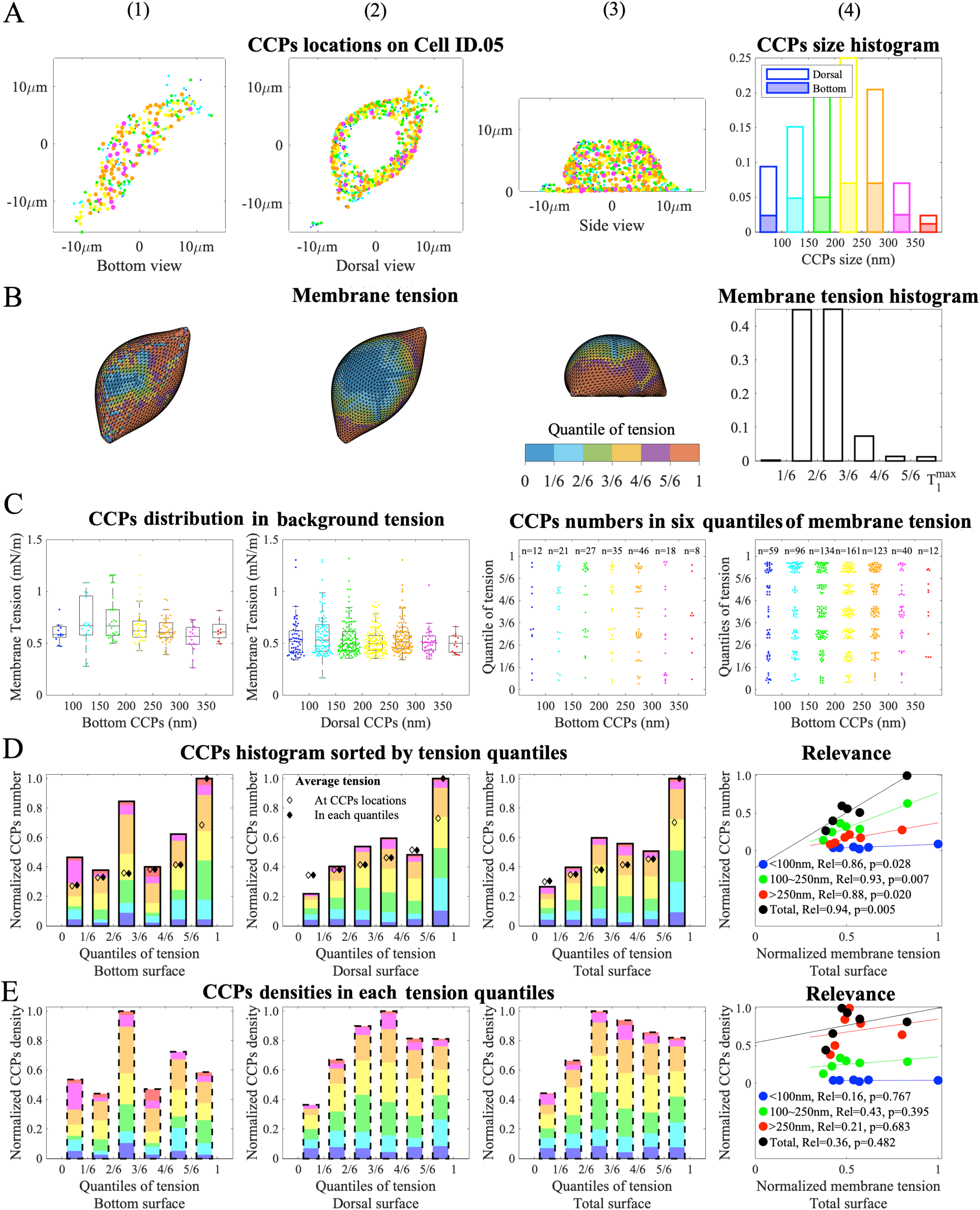
The comparison of the CCPs locations and the membrane tension on Cell ID.05.

**Figure S6.**
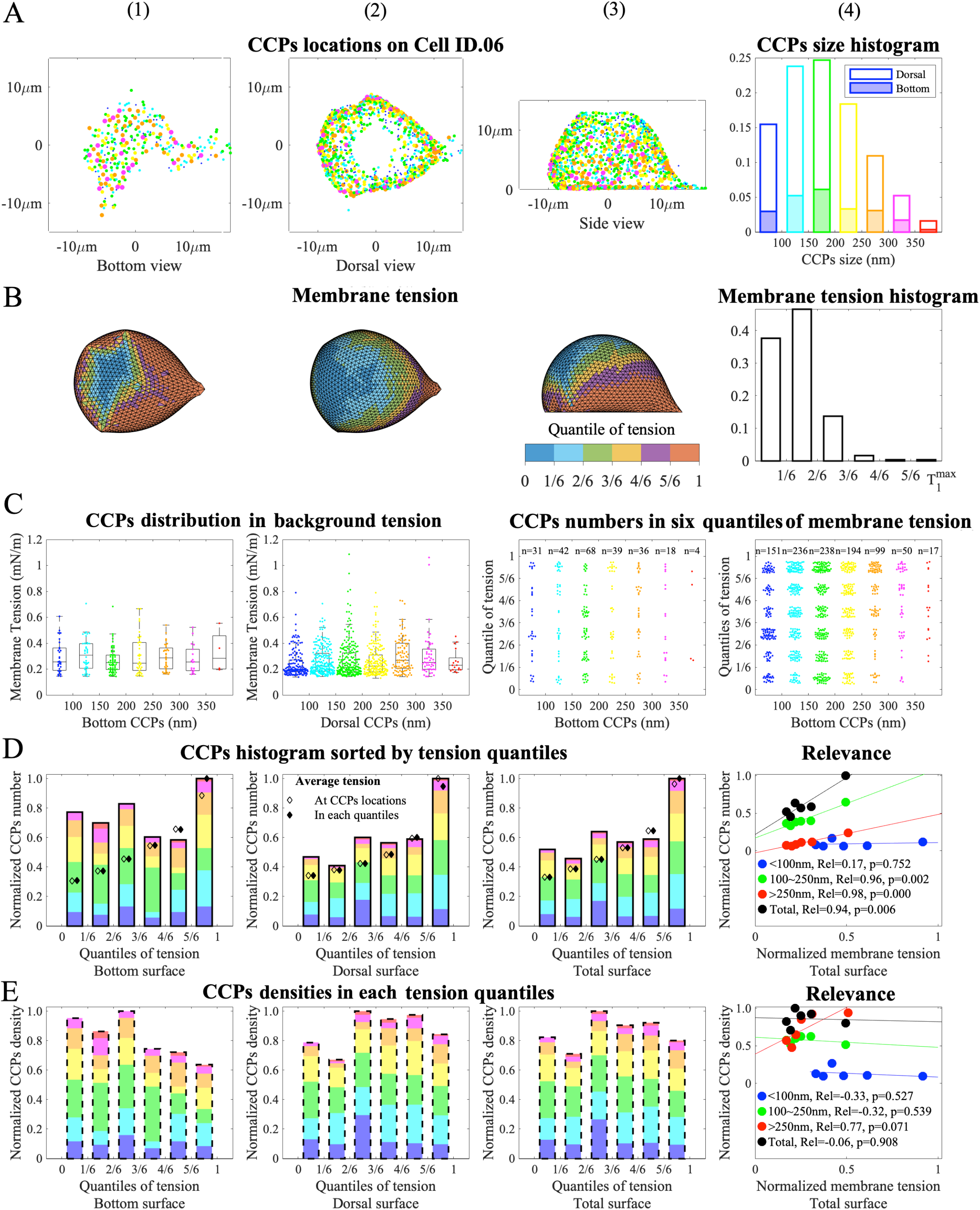
The comparison of the CCPs locations and the membrane tension on Cell ID.06.

**Figure S7.**
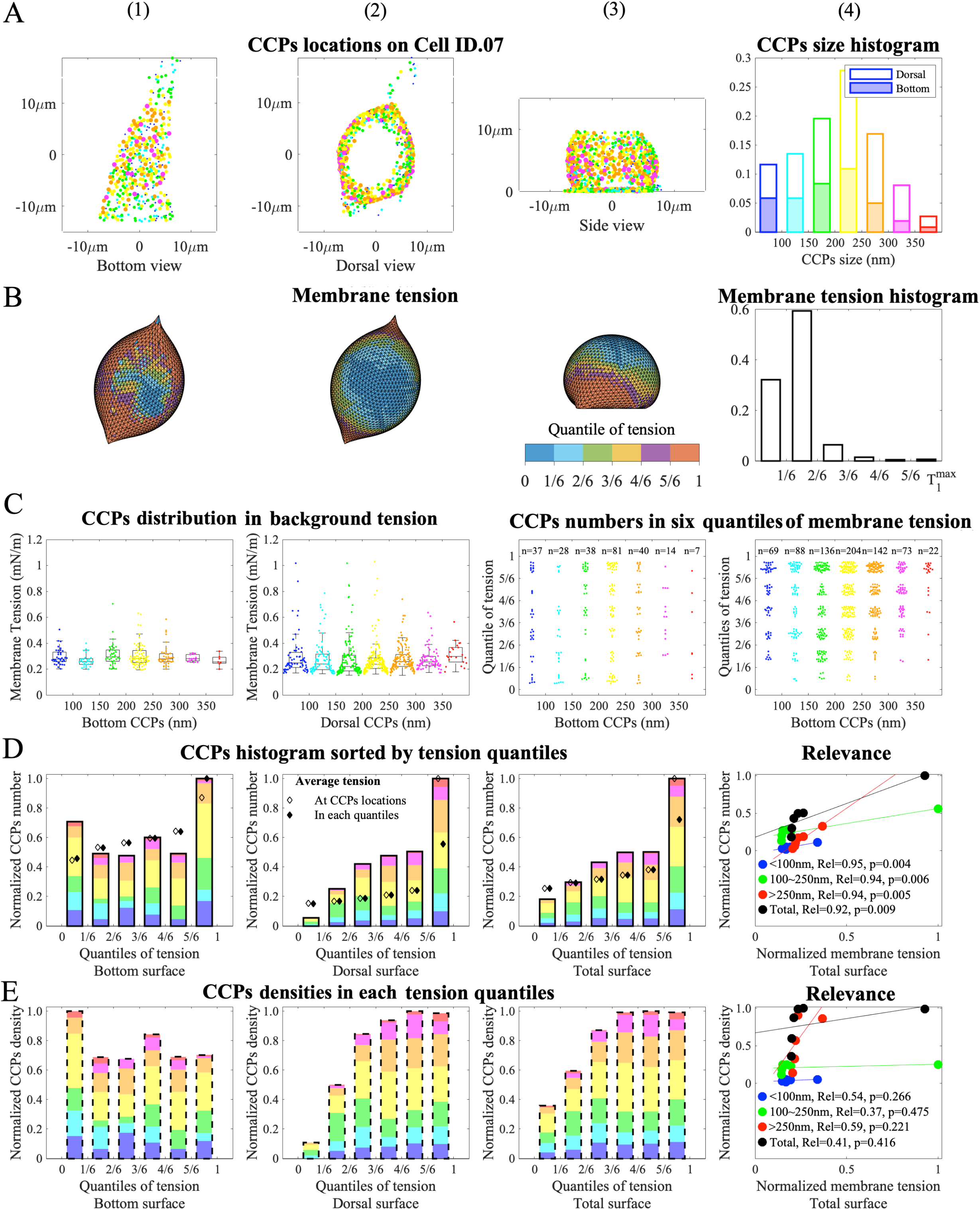
The comparison of the CCPs locations and the membrane tension on Cell ID.07.

**Figure S8.**
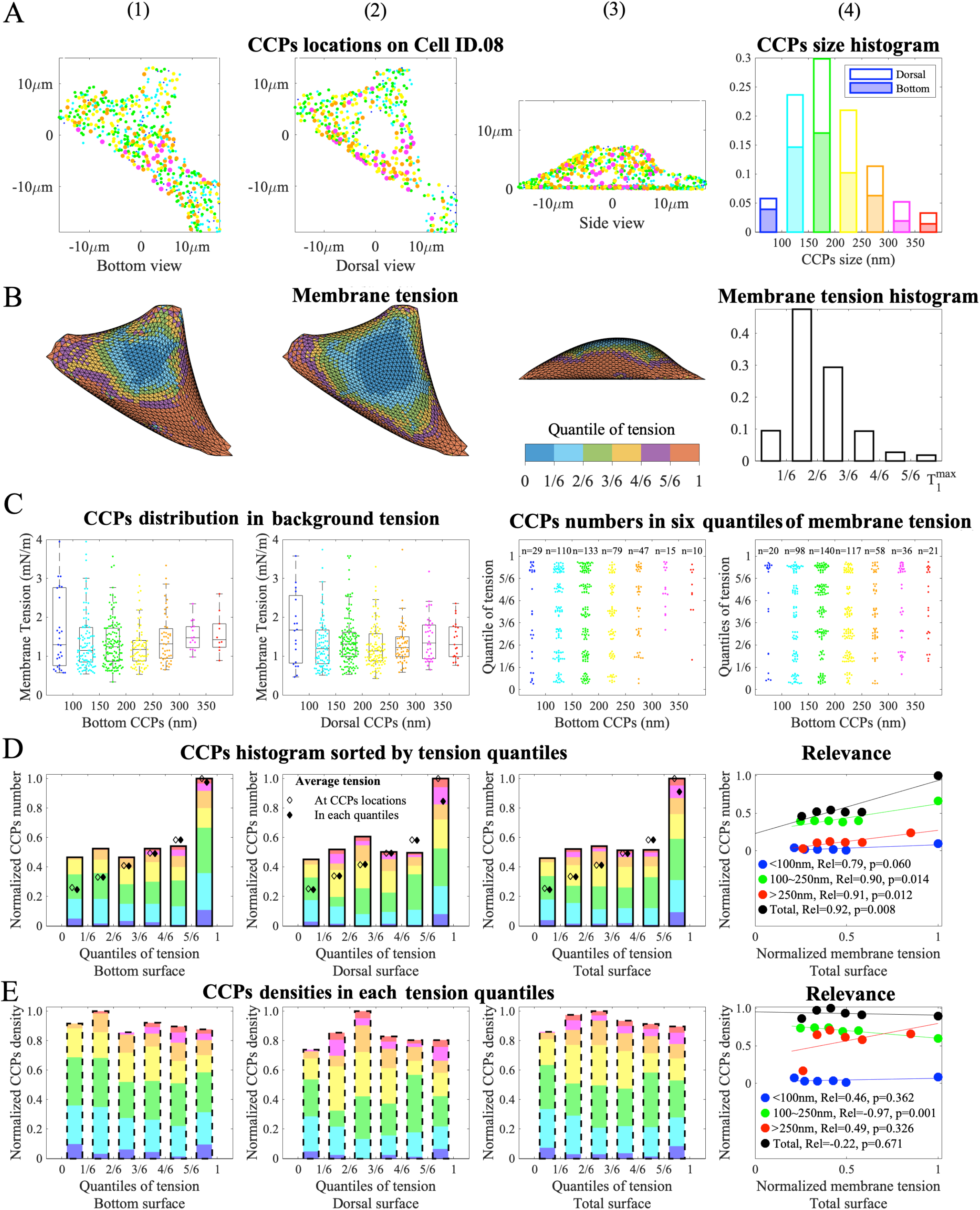
The comparison of the CCPs locations and the membrane tension on Cell ID.08.

**Figure S9.**
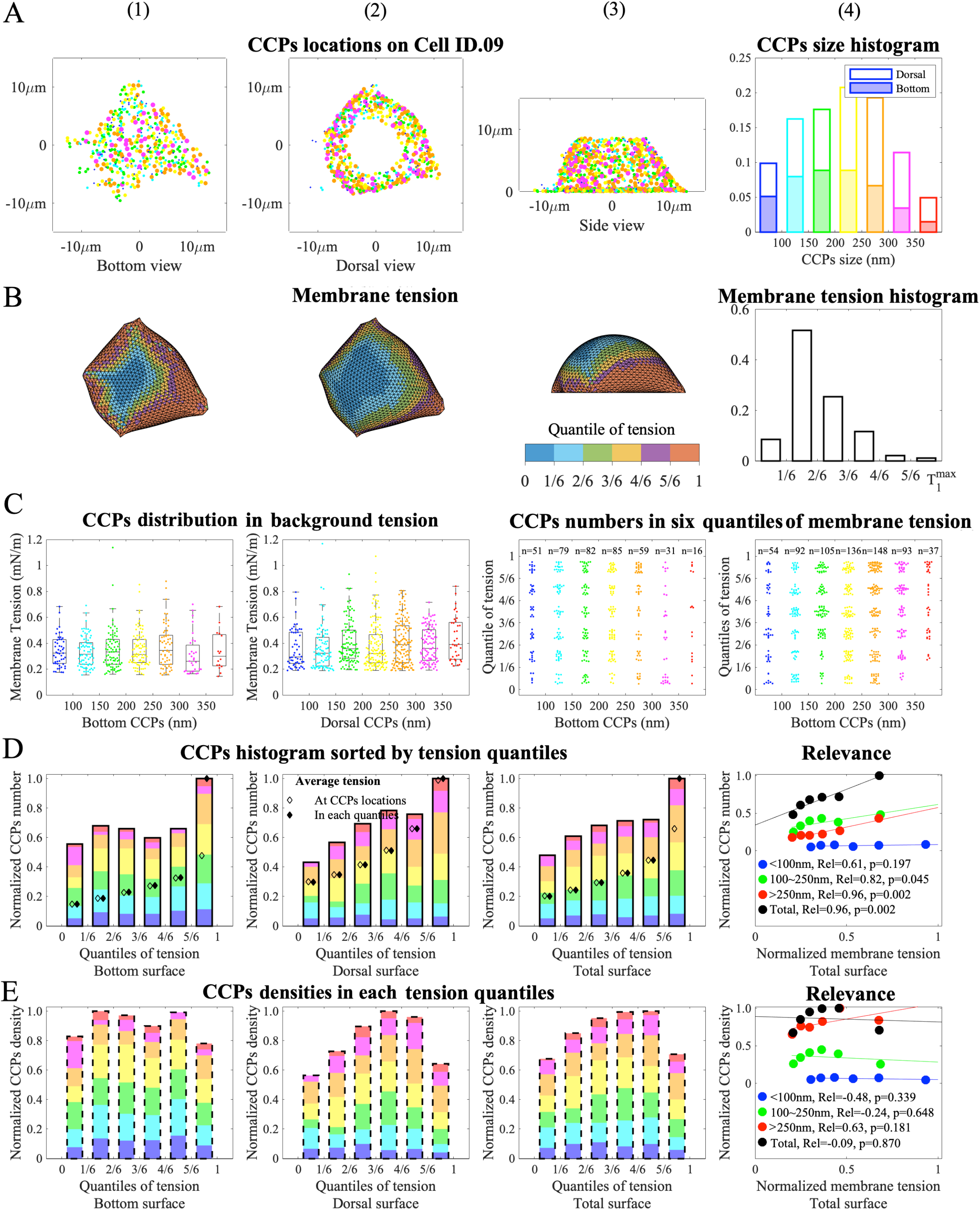
The comparison of the CCPs locations and the membrane tension on Cell ID.09.

**Figure S10.**
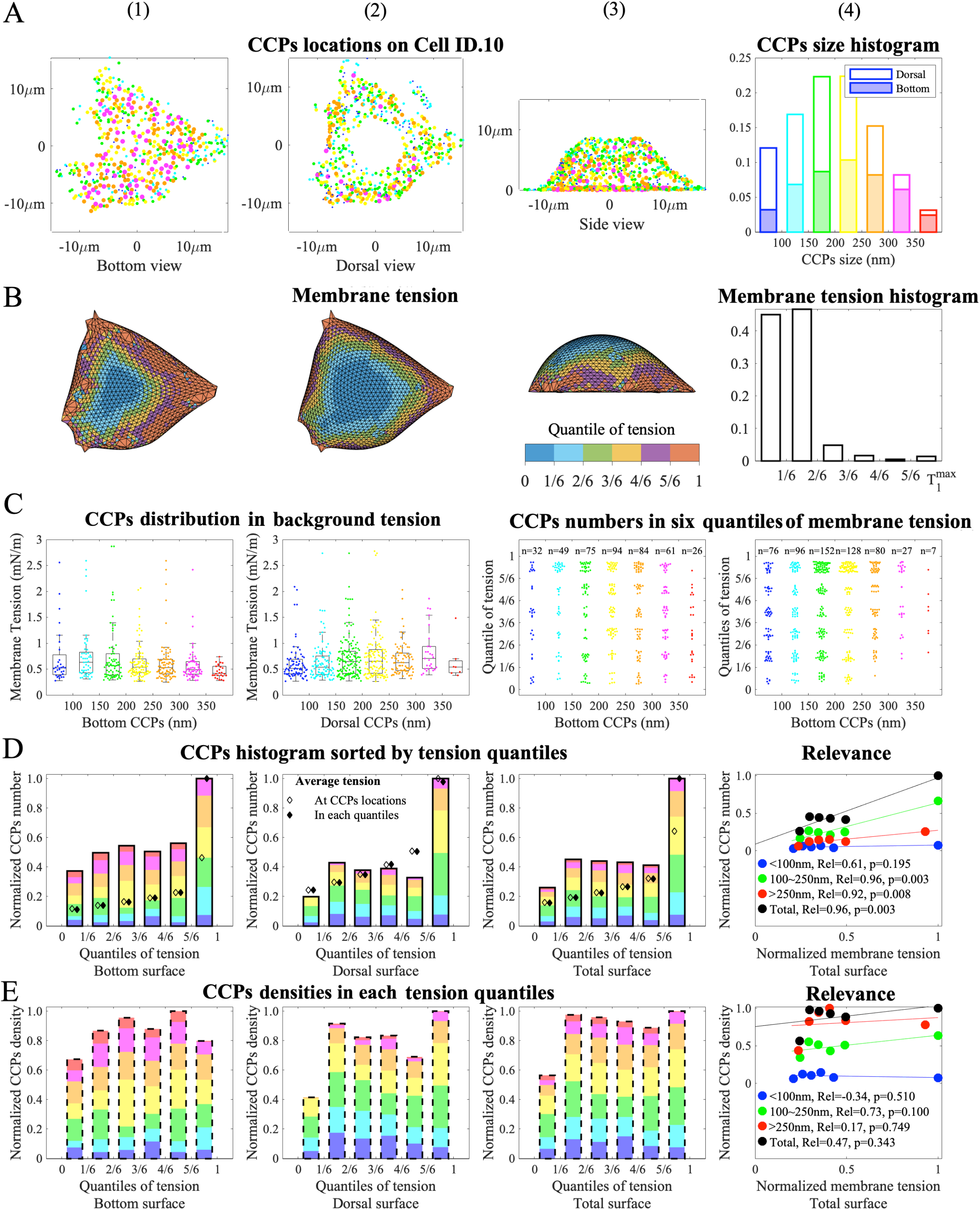
The comparison of the CCPs locations and the membrane tension on Cell ID.10.

**Figure S11.**
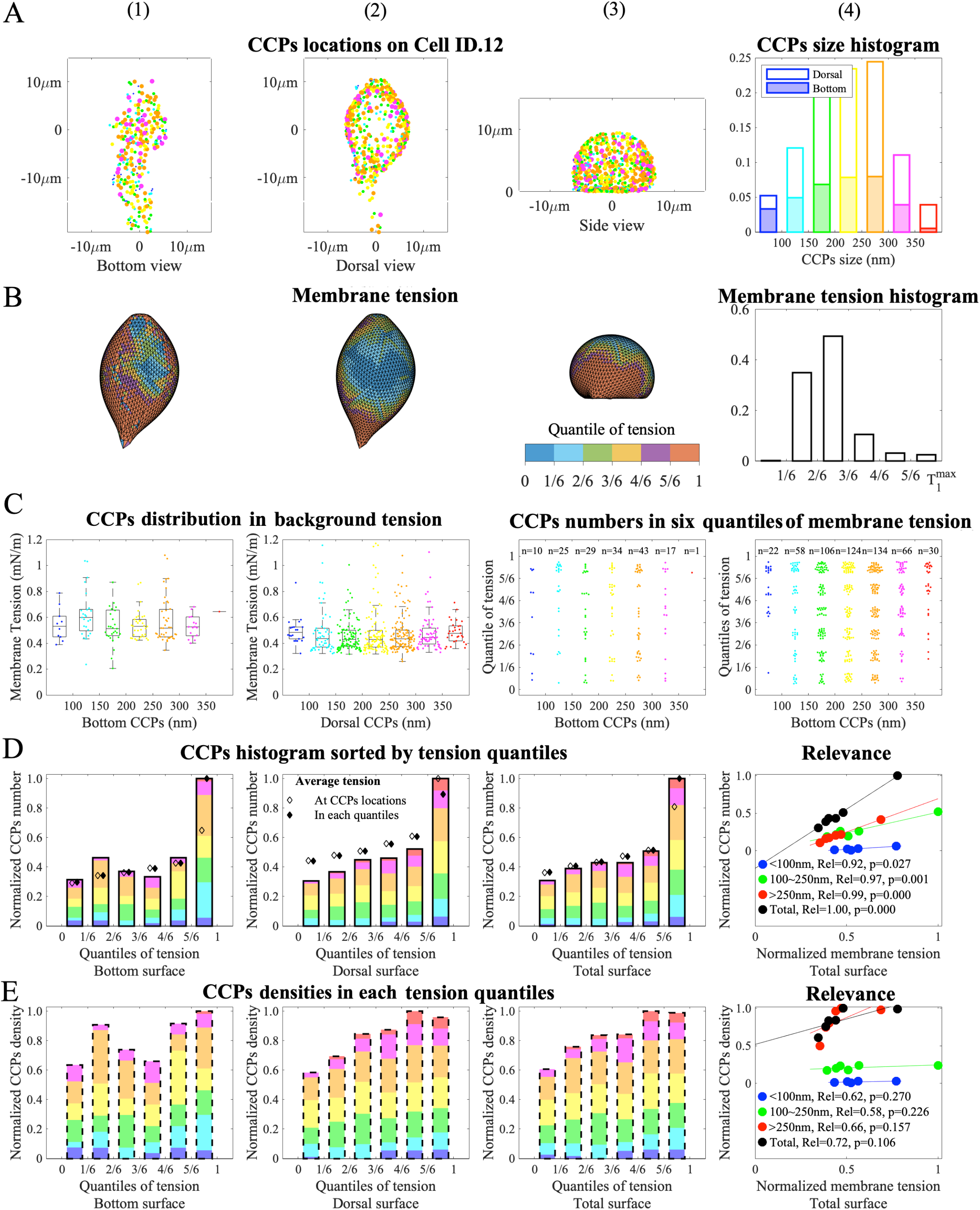
The comparison of the CCPs locations and the membrane tension on Cell ID.12.

**Table S1.**
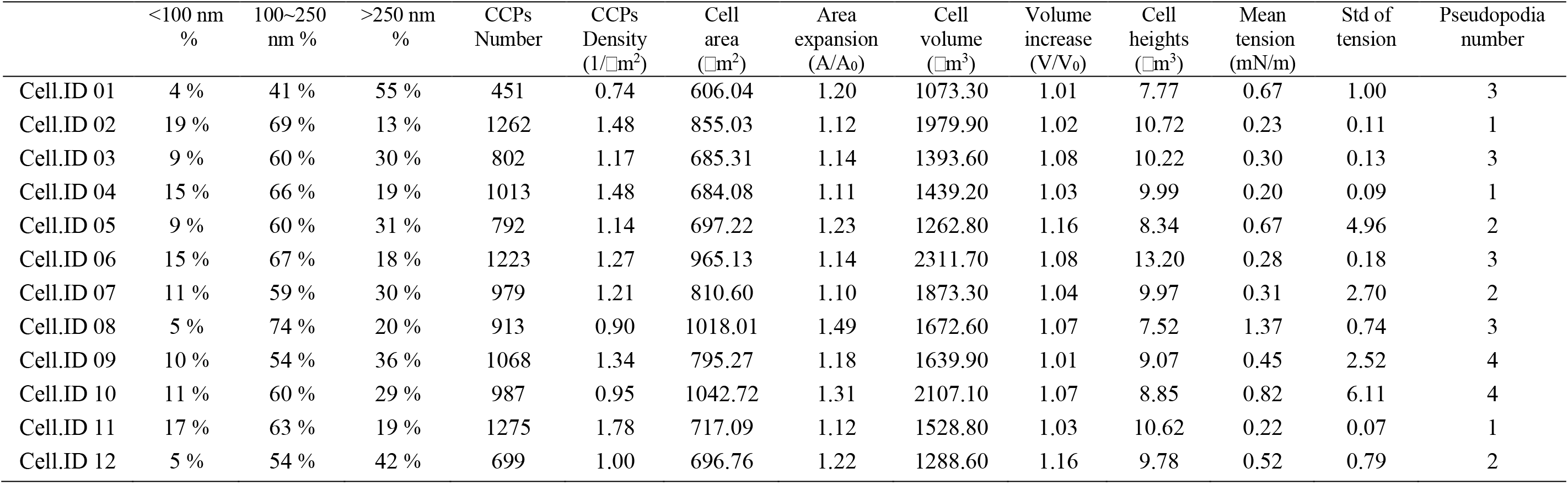
The values of parameters to describe the CCPs number, the cells geometry, and the membrane tension.

| | <100 nm<br>% | 100~250<br>nm % | >250 nm<br>% | CCPs<br>Number | CCPs<br>Density<br>(1/$\mu\text{m}^2$) | Cell<br>area<br>($\mu\text{m}^2$) | Area<br>expansion<br>(A/A<sub>0</sub>) | Cell<br>volume<br>($\mu\text{m}^3$) | Volume<br>increase<br>(V/V<sub>0</sub>) | Cell<br>heights<br>($\mu\text{m}^3$) | Mean<br>tension<br>(mN/m) | Std of<br>tension | Pseudopodia<br>number |
| --- | --- | --- | --- | --- | --- | --- | --- | --- | --- | --- | --- | --- | --- |
| Cell.ID 01 | 4 % | 41 % | 55 % | 451 | 0.74 | 606.04 | 1.20 | 1073.30 | 1.01 | 7.77 | 0.67 | 1.00 | 3 |
| Cell.ID 02 | 19 % | 69 % | 13 % | 1262 | 1.48 | 855.03 | 1.12 | 1979.90 | 1.02 | 10.72 | 0.23 | 0.11 | 1 |
| Cell.ID 03 | 9 % | 60 % | 30 % | 802 | 1.17 | 685.31 | 1.14 | 1393.60 | 1.08 | 10.22 | 0.30 | 0.13 | 3 |
| Cell.ID 04 | 15 % | 66 % | 19 % | 1013 | 1.48 | 684.08 | 1.11 | 1439.20 | 1.03 | 9.99 | 0.20 | 0.09 | 1 |
| Cell.ID 05 | 9 % | 60 % | 31 % | 792 | 1.14 | 697.22 | 1.23 | 1262.80 | 1.16 | 8.34 | 0.67 | 4.96 | 2 |
| Cell.ID 06 | 15 % | 67 % | 18 % | 1223 | 1.27 | 965.13 | 1.14 | 2311.70 | 1.08 | 13.20 | 0.28 | 0.18 | 3 |
| Cell.ID 07 | 11 % | 59 % | 30 % | 979 | 1.21 | 810.60 | 1.10 | 1873.30 | 1.04 | 9.97 | 0.31 | 2.70 | 2 |
| Cell.ID 08 | 5 % | 74 % | 20 % | 913 | 0.90 | 1018.01 | 1.49 | 1672.60 | 1.07 | 7.52 | 1.37 | 0.74 | 3 |
| Cell.ID 09 | 10 % | 54 % | 36 % | 1068 | 1.34 | 795.27 | 1.18 | 1639.90 | 1.01 | 9.07 | 0.45 | 2.52 | 4 |
| Cell.ID 10 | 11 % | 60 % | 29 % | 987 | 0.95 | 1042.72 | 1.31 | 2107.10 | 1.07 | 8.85 | 0.82 | 6.11 | 4 |
| Cell.ID 11 | 17 % | 63 % | 19 % | 1275 | 1.78 | 717.09 | 1.12 | 1528.80 | 1.03 | 10.62 | 0.22 | 0.07 | 1 |
| Cell.ID 12 | 5 % | 54 % | 42 % | 699 | 1.00 | 696.76 | 1.22 | 1288.60 | 1.16 | 9.78 | 0.52 | 0.79 | 2 |

## Notes

### Competing Interest Statement

The authors have declared no competing interest.

### Summary of Updates

Discussion revised, Figure 1 8 s1-11 revised, reference added

## References

[1] M. Kaksonen, A. Roux, Mechanisms of clathrin-mediated endocytosis, Nature reviews Molecular cell biology 19 (5) (2018) 313–326.

[2] S. Boulant, C. Kural, J.-C. Zeeh, F. Ubelmann, T. Kirchhausen, Actin dynamics counteract membrane tension during clathrin-mediated endocytosis, Nature cell biology 13 (9) (2011) 1124–1131.

[3] M. Saleem, S. Morlot, A. Hohendahl, J. Manzi, M. Lenz, A. Roux, A balance between membrane elasticity and polymerization energy sets the shape of spherical clathrin coats, Nature communications 6 (1) (2015) 6249.

[4] S. Morlot, V. Galli, M. Klein, N. Chiaruttini, J. Manzi, F. Humbert, L. Dinis, M. Lenz, G. Cappello, A. Roux, Membrane shape at the edge of the dynamin helix sets location and duration of the fission reaction, Cell 151 (3) (2012) 619–629.

[5] D. Loerke, M. Mettlen, D. Yarar, K. Jaqaman, H. Jaqaman, G. Danuser, S. L. Schmid, Cargo and dynamin regulate clathrin-coated pit maturation, PLoS biology 7 (3) (2009) e1000057.

[6] T. Kirchhausen, Imaging endocytic clathrin structures in living cells, Trends in cell biology 19 (11) (2009) 596–605.

[7] A. Banerjee, A. Berezhkovskii, R. Nossal, Stochastic model of clathrin-coated pit assembly, Biophysical journal 102 (12) (2012) 2725–2730.

[8] A. Banerjee, A. Berezhkovskii, R. Nossal, Distributions of lifetime and maximum size of abortive clathrin-coated pits, Physical Review E 86 (3) (2012) 031907.

[9] J. Heuser, Three-dimensional visualization of coated vesicle formation in fibroblasts., The Journal of cell biology 84 (3) (1980) 560–583.

[10] B. Pearse, R. Crowther, Structure and assembly of coated vesicles, Annual review of biophysics and biophysical chemistry 16 (1) (1987) 49–68.

[11] L. Schermelleh, R. Heintzmann, H. Leonhardt, A guide to super-resolution fluorescence microscopy, Journal of Cell Biology 190 (2) (2010) 165–175.

[12] D. Li, L. Shao, B.-C. Chen, X. Zhang, M. Zhang, B. Moses, D. E. Milkie, J. R. Beach, J. A. Hammer III, M. Pasham, et al., Extended-resolution structured illumination imaging of endocytic and cytoskeletal dynamics, Science 349 (6251) (2015) aab3500.

[13] N. M. Willy, J. P. Ferguson, A. Akatay, S. Huber, U. Djakbarova, S. Silahli, C. Cakez, F. Hasan, H. C. Chang, A. Travesset, et al., De novo endocytic clathrin coats develop curvature at early stages of their formation, Developmental cell 56 (22) (2021) 3146–3159.

[14] U. Djakbarova, Y. Madraki, E. T. Chan, C. Kural, Dynamic interplay between cell membrane tension and clathrin-mediated endocytosis, Biology of the Cell 113 (8) (2021) 344–373.

[15] J. Dai, H. P. Ting-Beall, M. P. Sheetz, The secretion-coupled endocytosis correlates with membrane tension changes in rbl 2h3 cells, The Journal of general physiology 110 (1) (1997) 1–10.

[16] J. P. Ferguson, N. M. Willy, S. P. Heidotting, S. D. Huber, M. J. Webber, C. Kural, Deciphering dynamics of clathrin-mediated endocytosis in a living organism, Journal of Cell Biology 214 (3) (2016) 347–358.

[17] N. C. Gauthier, T. A. Masters, M. P. Sheetz, Mechanical feedback between membrane tension and dynamics, Trends in cell biology 22 (10) (2012) 527–535.

[18] R. Yonashiro, A. Sugiura, M. Miyachi, T. Fukuda, N. Matsushita, R. Inatome, Y. Ogata, T. Suzuki, N. Dohmae, S. Yanagi, Mitochondrial ubiquitin ligase mitol ubiquitinates mutant sod1 and attenuates mutant sod1-induced reactive oxygen species generation, Molecular biology of the cell 20 (21) (2009) 4524–4530.

[19] X. Tan, J. Heureaux, A. P. Liu, Cell spreading area regulates clathrin-coated pit dynamics on micropatterned substrate, Integrative Biology 7 (9) (2015) 1033–1043.

[20] N. M. Willy, F. Colombo, S. Huber, A. C. Smith, E. G. Norton, C. Kural, E. Cocucci, Calm supports clathrin-coated vesicle completion upon membrane tension increase, Proceedings of the National Academy of Sciences 118 (25) (2021) e2010438118.

[21] N. Willy, J. Ferguson, S. Huber, S. Heidotting, E. Aygün, S. Wurm, E. Johnston-Halperin, M. Poirier, C. Kural, Membrane mechanics govern spatiotemporal heterogeneity of endocytic clathrin coat dynamics, Molecular biology of the cell 28 (24) (2017) 3480–3488.

[22] E. M. Batchelder, D. Yarar, Differential requirements for clathrin-dependent endocytosis at sites of cell–substrate adhesion, Molecular Biology of the Cell 21 (17) (2010) 3070–3079.

[23] A. A. Akatay, T. Wu, U. Djakbarova, C. Thompson, E. Cocucci, R. Zandi, J. Rudnick, C. Kural, Endocytosis at extremes: formation and internalization of giant clathrin-coated pits under elevated membrane tension, Frontiers in Molecular Biosciences 9 (2022) 959737.

[24] N. C. Gauthier, Plasma membrane area increases with spread area by exocytosis of gpi anchored protein compartment, Biophysical Journal 96 (3) (2009) 151a.

[25] E. Kreysing, J. M. Hugh, S. K. Foster, K. Andresen, R. D. Greenhalgh, E. K. Pillai, A. Dimitracopoulos, U. F. Keyser, K. Franze, Effective cell membrane tension is independent of polyacrylamide substrate stiffness, PNAS nexus 2 (1) (2023) pgac299.

[26] F. Tebar, S. K. Bohlander, A. Sorkin, Clathrin assembly lymphoid myeloid leukemia (calm) protein: localization in endocytic-coated pits, interactions with clathrin, and the impact of overexpression on clathrin-mediated traffic, Molecular biology of the cell 10 (8) (1999) 2687–2702.

[27] S. E. Miller, S. Mathiasen, N. A. Bright, F. Pierre, B. T. Kelly, N. Kladt, A. Schauss, C. J. Merrifield, D. Stamou, S. Höning, et al., Calm regulates clathrin-coated vesicle size and maturation by directly sensing and driving membrane curvature, Developmental cell 33 (2) (2015) 163–175.

[28] F. Aguet, C. N. Antonescu, M. Mettlen, S. L. Schmid, G. Danuser, Advances in analysis of low signal-to-noise images link dynamin and ap2 to the functions of an endocytic checkpoint, Developmental cell 26 (3) (2013) 279–291.

[29] J. Z. Rappoport, S. Kemal, A. Benmerah, S. M. Simon, Dynamics of clathrin and adaptor proteins during endocytosis, American Journal of Physiology-Cell Physiology 291 (5) (2006) C1072–C1081.

[30] M. Akamatsu, R. Vasan, D. Serwas, M. A. Ferrin, P. Rangamani, D. G. Drubin, Principles of self-organization and load adaptation by the actin cytoskeleton during clathrin-mediated endocytosis, Elife 9 (2020) e49840.

[31] M. Wu, X. Wu, A kinetic view of clathrin assembly and endocytic cargo sorting, Current opinion in cell biology 71 (2021) 130–138.

[32] K. A. Sochacki, B. L. Heine, G. J. Haber, J. R. Jimah, B. Prasai, M.A. Alfonzo-Méndez, A. D. Roberts, A. Somasundaram, J. E. Hinshaw, J. W. Taraska, The structure and spontaneous curvature of clathrin lattices at the plasma membrane, Developmental cell 56 (8) (2021) 1131–1146.

[33] W. F. Zeno, J. B. Hochfelder, A. S. Thatte, L. Wang, A. K. Gadok, C. C. Hayden, E. M. Lafer, J. C. Stachowiak, Clathrin senses membrane curvature, Biophysical Journal 120 (5) (2021) 818–828.

[34] G. Tagiltsev, C. A. Haselwandter, S. Scheuring, Nanodissected elastically loaded clathrin lattices relax to increased curvature, Science Advances 7 (33) (2021) eabg9934.

[35] D. K. Cureton, C. E. Harbison, E. Cocucci, C. R. Parrish, T. Kirchhausen, Limited transferrin receptor clustering allows rapid diffusion of canine parvovirus into clathrin endocytic structures, Journal of virology 86 (9) (2012) 5330–5340.

[36] M. Ehrlich, W. Boll, A. Van Oijen, R. Hariharan, K. Chandran, M. L. Nibert, T. Kirchhausen, Endocytosis by random initiation and stabilization of clathrin-coated pits, Cell 118 (5) (2004) 591–605.

[37] T. J. Nawara, A. L. Mattheyses, Imaging nanoscale axial dynamics at the basal plasma membrane, The International Journal of Biochemistry & Cell Biology 156 (2023) 106349.

[38] X. Liu, K.-c. Tsubota, Y. Yu, W. Xi, X. Gong, A numerical method to predict the membrane tension distribution of spreading cells based on the reconstruction of focal adhesions, Science China Physics, Mechanics & Astronomy 65 (6) (2022) 264612.

[39] X. Liu, H. Yang, Y. Liu, X. Gong, H. Huang, Numerical study of clathrin-mediated endocytosis of nanoparticles by cells under tension, Acta Mechanica Sinica 35 (2019) 691–701.

[40] M. G. Gustafsson, Nonlinear structured-illumination microscopy: wide-field fluorescence imaging with theoretically unlimited resolution, Proceedings of the National Academy of Sciences 102 (37) (2005) 13081–13086.

[41] P.-Y. Lin, J. Ge, C. Kuang, F.-J. Kao, Fluorescence detection and lifetime imaging with stimulated emission, Optical Nanoscopy and Novel Microscopy Techniques (2014) 161.

[42] D. Thomann, D. R. Rines, P. K. Sorger, G. Danuser, Automatic fluorescent tag detection in 3d with super-resolution: application to the analysis of chromosome movement, Journal of microscopy 208 (1) (2002) 49–64.

[43] J.-P. Grossier, G. Xouri, B. Goud, K. Schauer, Cell adhesion defines the topology of endocytosis and signaling, The EMBO journal 33 (1) (2014) 35–45.

[44] N. Resolution, Imaging intracellular fluorescent proteins at, Protein Sci 9 (2000) 10.

[45] K. Jaqaman, D. Loerke, M. Mettlen, H. Kuwata, S. Grinstein, S. L. Schmid, G. Danuser, Robust single-particle tracking in live-cell time-lapse sequences, Nature methods 5 (8) (2008) 695–702.

[46] Y. Yang, D. Xiong, A. Pipathsouk, O. D. Weiner, M. Wu, Clathrin assembly defines the onset and geometry of cortical patterning, Developmental cell 43 (4) (2017) 507–521.

[47] A. Grassart, A. T. Cheng, S. H. Hong, F. Zhang, N. Zenzer, Y. Feng, D. M. Briner, G. D. Davis, D. Malkov, D. G. Drubin, Actin and dynamin2 dynamics and interplay during clathrin-mediated endocytosis, Journal of Cell Biology 205 (5) (2014) 721–735.

[48] A. J. Jin, K. Prasad, P. D. Smith, E. M. Lafer, R. Nossal, Measuring the elasticity of clathrin-coated vesicles via atomic force microscopy, Biophysical journal 90 (9) (2006) 3333–3344.

[49] G. Wang, T. Galli, Reciprocal link between cell biomechanics and exocytosis, Traffic 19 (10) (2018) 741–749.

[50] A. Diz-Muñoz, D. A. Fletcher, O. D. Weiner, Use the force: membrane tension as an organizer of cell shape and motility, Trends in cell biology 23 (2) (2013) 47–53.

[51] K. Keren, Cell motility: the integrating role of the plasma membrane, European Biophysics Journal 40 (2011) 1013–1027.

[52] A. Pietuch, A. Janshoff, Mechanics of spreading cells probed by atomic force microscopy, Open biology 3 (7) (2013) 130084.

[53] D. Raucher, M. P. Sheetz, Cell spreading and lamellipodial extension rate is regulated by membrane tension, The Journal of cell biology 148 (1) (2000) 127–136.

[54] A. R. Houk, A. Jilkine, C. O. Mejean, R. Boltyanskiy, E. R. Dufresne, S. B. Angenent, S. J. Altschuler, L. F. Wu, O. D. Weiner, Membrane tension maintains cell polarity by confining signals to the leading edge during neutrophil migration, Cell 148 (1) (2012) 175–188.

[55] A. D. Lieber, S. Yehudai-Resheff, E. L. Barnhart, J. A. Theriot, K. Keren, Membrane tension in rapidly moving cells is determined by cytoskeletal forces, Current biology 23 (15) (2013) 1409–1417.

[56] B. Huang, W. Wang, M. Bates, X. Zhuang, Three-dimensional super-resolution imaging by stochastic optical reconstruction microscopy, Science 319 (5864) (2008) 810–813.

[57] F. Aguet, S. Upadhyayula, R. Gaudin, Y.-y. Chou, E. Cocucci, K. He, B.-C. Chen, K. Mosaliganti, M. Pasham, W. Skillern, et al., Membrane dynamics of dividing cells imaged by lattice light-sheet microscopy, Molecular biology of the cell 27 (22) (2016) 3418–3435.

[58] D. Bucher, F. Frey, K. A. Sochacki, S. Kummer, J.-P. Bergeest, W. J. Godinez, H.-G. Kräusslich, K. Rohr, J. W. Taraska, U. S. Schwarz, et al., Clathrin-adaptor ratio and membrane tension regulate the flat-to-curved transition of the clathrin coat during endocytosis, Nature communications 9 (1) (2018) 1109.

[59] B. L. Scott, K. A. Sochacki, S. T. Low-Nam, E. M. Bailey, Q. Luu, A. Hor, A. M. Dickey, S. Smith, J. G. Kerkvliet, J. W. Taraska, et al., Membrane bending occurs at all stages of clathrin-coat assembly and defines endocytic dynamics, Nature communications 9 (1) (2018) 419.

[60] K. A. Sochacki, A. M. Dickey, M.-P. Strub, J. W. Taraska, Endocytic proteins are partitioned at the edge of the clathrin lattice in mammalian cells, Nature cell biology 19 (4) (2017) 352–361.

[61] M. G. Ford, I. G. Mills, B. J. Peter, Y. Vallis, G. J. Praefcke, P. R. Evans, H. T. McMahon, Curvature of clathrin-coated pits driven by epsin, Nature 419 (6905) (2002) 361–366.

[62] H. T. McMahon, J. L. Gallop, Membrane curvature and mechanisms of dynamic cell membrane remodelling, Nature 438 (7068) (2005) 590–596.

